# Localized molecular chaperone synthesis maintains neuronal dendrite proteostasis

**DOI:** 10.1101/2023.10.03.560761

**Authors:** Célia Alecki, Javeria Rizwan, Phuong Le, Suleima Jacob-Tomas, Mario Fernandez-Comaduran, Morgane Verbrugghe, Jia Ming Stella Xu, Sandra Minotti, James Lynch, Tad Wu, Heather Durham, Gene W. Yeo, Maria Vera

**Affiliations:** Department of Biochemistry, McGill University, Montreal, Quebec H3G 1Y6, Canada; Department of Cellular and Molecular Medicine, University of California, San Diego, La Jolla, CA, USA; Department of Neurology and Neurosurgery and Montreal Neurological Institute, McGill University, Montreal, Quebec H3A 2B4, Canada

**Author notes:** Department of Neurology and Neurosurgery, McGill University, Montreal, Canada (Suleima Jacob-Tomas). Department of Pathology, McGill University, Montreal, Canada (Tad Wu).

**Keywords:** Neuron, mRNA, protein homeostasis, dendrite, localized translation, neurodegeneration, heat shock protein, single-molecule fluorescence microscopy, amyotrophic lateral sclerosis

## Abstract

Proteostasis is maintained through regulated protein synthesis and degradation and chaperone-assisted protein folding. However, this is challenging in neuronal projections because of their polarized morphology and constant synaptic proteome remodeling. Using high-resolution fluorescence microscopy, we discovered that hippocampal and spinal cord motor neurons of mouse and human origin localize a subset of chaperone mRNAs to their dendrites and use microtubule-based transport to increase this asymmetric localization following proteotoxic stress. The most abundant dendritic chaperone mRNA encodes a constitutive heat shock protein 70 family member (HSPA8). Proteotoxic stress also enhanced *HSPA8* mRNA translation efficiency in dendrites. Stress-mediated *HSPA8* mRNA localization to the dendrites was impaired by depleting fused in sarcoma—an amyotrophic lateral sclerosis-related protein—in cultured spinal cord mouse motor neurons or by expressing a pathogenic variant of heterogenous nuclear ribonucleoprotein A2/B1 in neurons derived from human induced pluripotent stem cells. These results reveal a crucial and unexpected neuronal stress response in which RNA-binding proteins increase the dendritic localization of *HSPA8* mRNA to maintain proteostasis and prevent neurodegeneration.

## Introduction

Cells have developed intricate mechanisms to maintain proteostasis—that is, to ensure that proteins are synthesized, folded, and degraded as needed. This is particularly challenging for neurons, because their complex polarized morphology includes projections that can span long distances and require constant proteome adjustments to respond to neuronal stimuli ^1–7^. Neuronal activity remodels the axonal terminal proteome ^8–10^. Likewise, stimulating individual dendritic spines triggers unique proteome changes independently from other spines ^11–14^. Neurons remodel local proteomes through the targeted distribution and regulation of the protein synthesis and degradation machinery (*e.g*., the proteasome and autophagy system to degrade damaged and unnecessary proteins) ^15–20^. Neurons ensure timely and efficient protein synthesis through an at-a-distance expression mechanism that relies on localizing specific mRNAs and regulating their stability and translation ^12,21–26^. Thus, neurons tightly regulate the distribution of proteins enriched in axons, dendrites, and synapses by localizing their mRNAs and necessary translation factors to these regions while retaining other mRNAs in the soma ^24,27–30^. Axon- and dendrite-targeted mRNAs contain specific sequence/structure motifs (zip codes) recognized by particular RNA binding proteins (RBPs) ^31,32^. Selective interactions between RBPs and motors form unique complexes or neuronal granules ^33–38^, which move mRNAs in both directions by anchoring them to microtubule motors (dynein and kinesins) or membranous organelles for active transport to axons and dendrites ^24,27,39,40^. Some RBPs also prevent mRNA translation during transport and derepress translation in response to local synaptic stimuli ^41–43^.

Successful protein synthesis and targeted degradation require the chaperoning function of heat shock proteins (HSPs). HSPs facilitate the folding of newly synthesized polypeptides into their functional three-dimensional conformations, and subsequently sequester or refold proteins that take on abnormal conformations, preventing aggregation and aberrant interactions in the crowded intracellular environment ^44–47^. Multiple HSPs load onto a misfolded substrate and perform several refolding cycles to restore proper conformation and sustain proteostasis ^48^. HSPs are grouped into families based on their molecular weights ^45,49^. The HSP60, HSP70, and HSP90 families actively promote protein folding in all cell types ^50–52^, and their functions are modulated by diverse co-chaperones ^53–56^. They also cooperate with small HSPs (sHSPs) ^52,53,57^ and HSP110 (HSPH) ^58^ to prevent and resolve misfolded protein aggregation and target misfolded proteins for degradation ^59–62^. Accordingly, subsets of them localize prominently to the dendrites and axons in diverse neuronal types ^63–66^. Intriguingly, neurons subjected to certain proteotoxic stresses have elevated levels of HSP70 and DNAJ (HSP40) family members in their dendrites and synapses ^63,67–69^.

The mechanism underlying HSP subcellular distribution in neurons represents a major knowledge gap. Most studies on induced HSP expression have investigated their upregulation by the transcription factor heat shock factor 1 (HSF1) ^47,70–73^. Recently, mRNAs encoding HSPA8 and HSP90AA were identified in the dendritic transcriptome under basal conditions ^74^. Here, we report that neurons increase the transport and local translation of a subset of HSP mRNAs in the dendrites in response to proteotoxic stress. Combining high-resolution fluorescence microscopy and molecular biology, we characterized changes in the subcellular localization of HSP mRNAs in primary hippocampal and spinal cord motor neurons subjected to different proteotoxic insults. Fused in sarcoma (FUS) and heterogenous nuclear ribonucleoprotein A2/B1 (HNRNPA2B1), both implicated in amyotrophic lateral sclerosis (ALS), were identified as important regulators of the subcellular distribution of the constitutive HSP70 mRNA, HSPA8, and their actions were essential for dendritic proteostasis during stress.

## Results

### Hippocampal neurons alter HSP mRNA distributions upon stress

To study how neurons tailor HSP expression to proteostatic demands, we subjected cultures of primary mouse hippocampal neurons to proteotoxic stress by inhibiting the proteasome. These cultures faithfully recapitulate the regulation of mRNA localization and translation in response to neuronal stimuli and the activation of HSP transcription upon stress ^12,75–78^. Dissociated cultures of hippocampi from postnatal day 0 mouse pups differentiate to express features of their mature *in situ* counterparts by day 17. Treatment with the proteasome inhibitor MG132 (10 μM for 7 h) results in the accumulation of misfolded proteins and protein quality failure—prominent hallmarks of neurodegenerative disorders ^79^. To study MG132-induced changes in subcellular mRNA localization, we isolated the total RNA from somas and projections separately harvested from neurons cultured in Transwell membrane filter inserts ^80^ (**Fig. 1a**). RNA sequencing (RNA-Seq) and differential expression analysis (DESeq2) revealed previously described transcript signatures specific to somas and projections in steady-state (control (Ctrl)) neurons, *e.g*., dendritic localization of calcium/calmodulin-dependent protein kinase II alpha (*CamK2a*) and β-*actin*) ^74,81^. Exposure to MG132 (10 μM for 7 h) significantly changed the expression of hundreds of RNAs between the soma and neurites (**Fig. 1b**). In fact, in gene ontology analyses of the changed transcripts, the only biological function enriched in both compartments was “protein refolding” (**Fig. 1c, S1a– S1c and Table S1**). Tens of transcripts coding for constitutive protein folding chaperones, such as *Hsp90ab*, *Hspa8*, and *Hsp110*, were previously identified in the soma and projections of hippocampal neurons obtained from rodent tissue and cultured primary neurons under homeostatic conditions ^11,74,82^, but changes in their subcellular distribution in response to proteotoxic stress have not been analyzed.

**Fig. 1.**
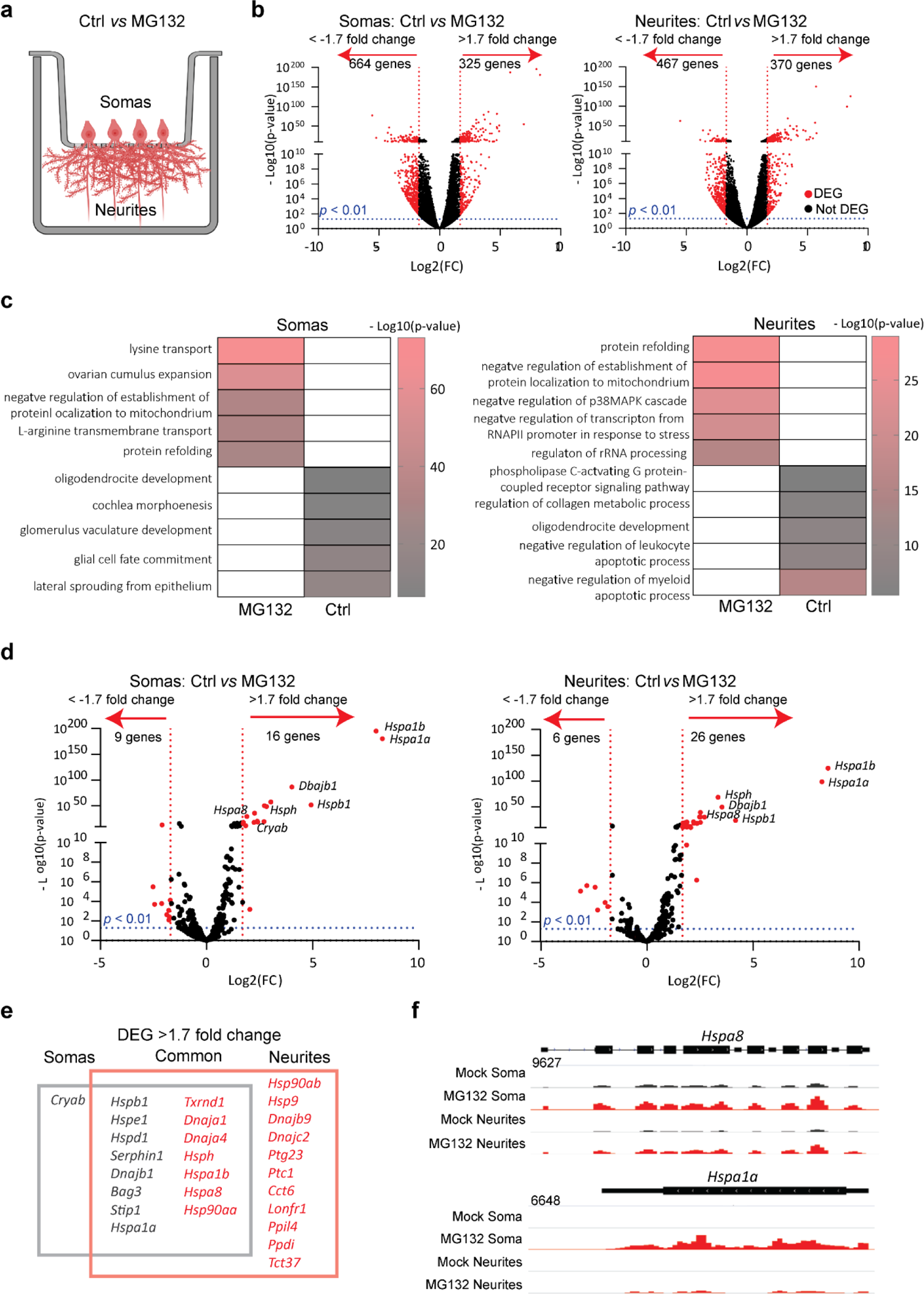
Specific mRNAs are preferentially enriched in the soma or projections of hippocampal neurons after proteostatic stress. **(a)** Schematic of primary mouse hippocampal neurons cultured in Transwell membranes to physically separate the soma and neurites for RNA extraction. Neurons were exposed to MG132 (10 μM for 7 h) or DMSO (Ctrl). **(b)** Volcano plot of differentially expressed genes (DEGs) in the soma or neurites (*n* = 3). Genes up- or down-regulated by > 1.7 fold after MG132 treatment and *P*-values < 0.01 are indicated in red. **(c)** Gene ontology enrichment analysis. Gene ontology categories of the top five biological processes enriched in DEGs in the somas and neurites after MG132 exposure. The color of the bands denotes the extent of upregulation. **(d)** Volcano plot of known chaperone-related genes. Genes up- or down-regulated by > 1.7 fold after MG132 treatment and *P*-values < 0.01 are indicated in red. **(e)** Venn diagram listing the differentially enriched molecular chaperone-related genes in the somas (gray square) and neurites (red square). (**f)** RNA-seq distributions of the *Hspa8* and *Hspa1a1* loci in the soma and neurites of control and MG132-exposed neurons.

Mammals contain over 400 genes encoding molecular chaperones and co-chaperones ^83^; of these, only 16 were upregulated in both fractions with increased enrichment in either the soma (*e.g*., the inducible HSP70 *Hspa1a*) or the projections (*e.g*., the constitutive HSP70 *Hspa8*). Interestingly, while mRNAs for 11 chaperones were specifically increased in neuronal projections, only the mRNA encoding the sHSP CRYAB was enriched in the soma (**Fig. 1d–1f**). Importantly, co-chaperones colocalized with their chaperone partners and HSP mRNA distributions matched the subcellular locations of their known folding clients. For instance, *Hsp90aa* and *Hsp90ab* and their co-chaperone *Ptg23* were enriched in projections upon stress. In dendrites, HSP90 supports the delivery of α-amino-3-hydroxy-5-methyl4-isoxazolepropionic acid receptors to the spine membrane, which is critical for synaptic transmission in the hippocampus ^84^. Likewise, DNAJs localized with their refolding partners, constitutive (HSPA8) and inducible (HSPA1A) HSP70s. In contrast with the significant upregulation of HSP mRNAs in projections, the levels of *Camk2a* and β-*actin* mRNAs, which are well-known to localize to dendrites, were unchanged ^74,81^ (**Fig. S1d**). These data suggest that neurons identify the need for HSP and co-chaperone mRNAs and distribute them to the same compartments as their client proteins.

### A subset of HSP mRNAs specifically localize to the dendrites upon stress

To define the principles of selective neuronal HSP mRNA localization and to identify proteotoxicity-induced changes in their subcellular distributions, we combined single-molecule fluorescence *in situ* hybridization (smFISH) with immunofluorescence (IF) to localize single mRNAs in primary hippocampal neurons using established markers of dendrites (microtubule-associated protein 2 (MAP2)), axons (microtubule-associated protein tau (TAU)), and spines (postsynaptic density 95 (PSD95); **Fig. 2a**) ^85,86^. Single mRNAs and transcription sites were identified with validated specific probes (**Fig. S2a and S2b**). We used the computational pipeline FISH-quant and its point spread function (PSF) superimposition approach to quantify single mRNAs and nascent transcripts, respectively ^87,88^. To collect statistics on mRNA localization across the neuronal morphology, we updated the computational pipeline Analysis of RNA Localization In Neurons (ARLIN) ^89^. This pipeline was used to validate the RNA-seq data by studying the main HSPs implicated in proteostasis loss during neurodegeneration: HSPA1A, HSPA8, HSP90AA, HSP90AB, and HSP110 ^50,58,66,90,91^. We first analyzed changes in the soma and quantified the significant transcriptional induction and increased concentrations of these constitutive and inducible HSP mRNAs, while the induction of *Dnajb1* and *Dnajb5* (used as controls) was lower; the same pattern was detected by RNA-seq (**Fig. 2b–d)**.

**Fig. 2.**
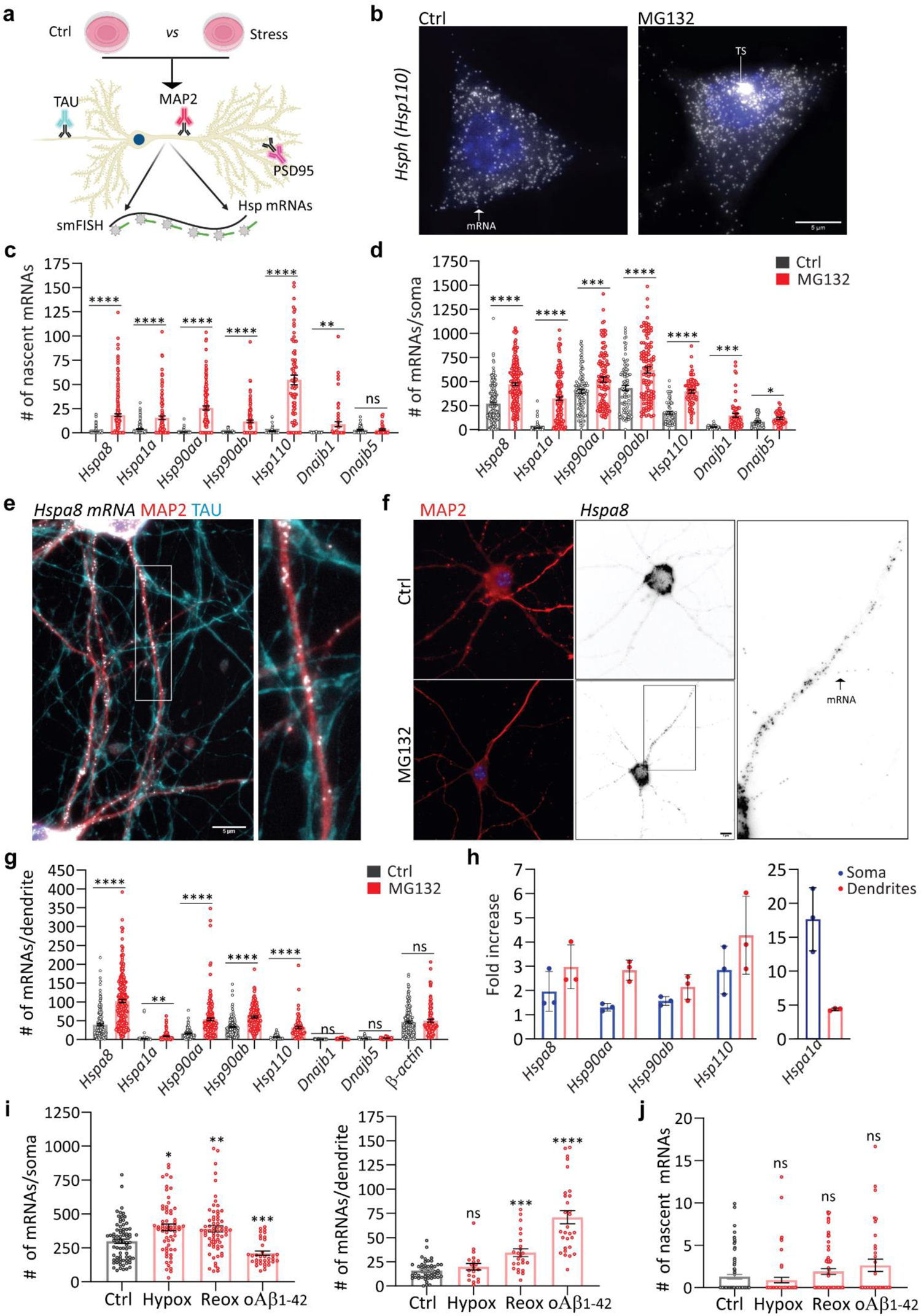
Subcellular distributions of HSP mRNAs in hippocampal neurons upon stress. **(a)** Schematic of the combined immunofluorescence (IF) and single-molecule fluorescence *in situ* hybridization (smFISH) protocol used on fixed primary hippocampal neurons. (**b**) smFISH detection of *Hsp110* mRNAs in the soma and nucleus (blue) of control (Ctrl) and MG132-stressed neurons. Arrows in the Ctrl and MG132 images indicate a single mRNA and a transcription site (TS), respectively. (**c, d)** Quantification of nascent transcripts (**C**) and somatic (**D**) HSP mRNAs in Ctrl and MG132-stressed neurons. Data are the mean ± standard error of the mean (SEM) of three independent experiments (*n* = 37-302 neurons total; dots indicate individual values). **(e)** Localization of *Hspa8* mRNA (smFISH, white) in the dendrites (IF: MAP2, red) and axons (IF: TAU, blue) of hippocampal neurons. Scale bar = 5 μm. The square depicts the magnified region. **(f)** Detection of *Hspa8* mRNA (smFISH, black) in dendrites (IF: MAP2, red) in Ctrl and MG132 stressed neurons. The square depicts the magnified region. **(g)** Quantification of dendritic HSP mRNAs in the Ctrl and MG132-stressed neurons in C, D (*n* = 45-248 dendrites). (**h)** Fold enrichment of HSP mRNAs in the soma and dendrites of MG132-stressed (MG) and Ctrl neurons from the quantifications in C, D, and G. (**i**) Quantification of somatic and dendritic *Hspa8* mRNAs in Ctrl hippocampal neurons and those stressed by hypoxia (Hypox), hypoxia followed by reoxygenation (Reox), or incubation with amyloid beta (1–42) oligomers (oAβ_1-42_) (*n* = 35-76 neurons and *n* = 24-48 dendrites). (**j**) Quantification of nascent *Hspa8* mRNA in one replicate of (I). Data are the mean ± SEM of two independent experiments (*n* = 35–77 neurons; dots indicate individual values). ****, *P* < 0.0001; ***, *P* < 0.001; **, *P* < 0.01; *, *P* < 0.05; ns, not significant (by unpaired *t*-test).

We next ascribed the mRNAs enriched in neuronal projections to either the axons or dendrites by combining smFISH with IF of TAU and MAP2. Remarkably, we did not detect any HSP mRNAs in the axons of Ctrl or MG132-stressed neurons (**Fig. 2e**). Instead, the mRNAs of all HSP of interest localized to the dendrites and were significantly enriched by MG132 (10 μM for 7 h) exposure (**Fig. 2f and 2g**). However, their concentrations varied greatly; from an average of 100 *Hspa8* mRNAs to only five *Hspa1a* mRNAs per dendrite. *Hspa1a* mRNA was retained in the soma, whereas *Hspa8*, *Hsp90aa*, and *Hsp110* mRNAs were enriched in the dendrites after MG132 stress, confirming the RNA-seq data (**Fig. 2h**). These results, together with the unchanging distribution of *β-actin* mRNA in the dendrites in response to MG132, strongly suggest that neurons selectively target specific HSP mRNAs to the dendrites upon stress (**Fig. 2g and 2h**).

Since *Hspa8* was the most abundant dendritic HSP mRNA measured, we verified that its dendritic localization occurs under disease-related stress conditions (**Fig. 2i and 2j**). We used hypoxia and hypoxia-reoxygenation injury (1% O_2_ for 3 h and 4 h reoxygenation), which generates reactive oxygen species resulting in protein misfolding ^92^ and brain damage ^93,94^ or neuronal exposure to 500 nM of oligomeric amyloid-β peptides for 24 h (oAβ_1-42_) ^95,96^, which accumulate in the hippocampus in Alzheimer’s disease and cause dendritic attrition ^95,96^. Dendritic *Hspa8* mRNA localization significantly increased after reoxygenation or oAβ_1-42_ exposure (**Fig. 2i**). Notably, although the level of transcriptional induction was much lower than after MG132 (10 μM for 7 h) exposure, these stresses relocated somatic *Hspa8* mRNAs to the dendrites (**Fig. 2i and 2j**). Therefore, the subcellular localization of *Hspa8* mRNAs to the dendrites is promoted by diverse proteotoxic stresses. These results suggest a common stress-induced regulatory mechanism that directs specific HSP mRNAs to the dendrites.

### Stress promotes HSP mRNA localization to the proximal dendrites and spines

We next investigated whether stress induces changes in the distribution of HSP mRNAs within the dendritic compartment, and particularly in the dendritic spines, which receive synaptic inputs. First, we considered that stress might drive HSP mRNAs into distal dendrites, broadening their distribution. To test this hypothesis, we measured the distance that each mRNA traveled from the soma and grouped them into bins of 25 μm to analyze their dendritic distributions (**Fig. 3a**). Stress induced significant increases in the number of *Hspa8*, *Hsp90aa*, *Hspa90ab*, and *Hsp110* mRNAs in the bins proximal to the soma. However, not significant increases in any of the mRNAs were observed at distances ≥ 125 μm from the soma. Thus, although the dendritic concentration of HSP mRNAs increased upon MG132-stress, their distribution over the dendritic shaft was comparable to that in non-stressed neurons, with more mRNAs in the proximal dendrites (**Fig. 3a**).

**Fig. 3.**
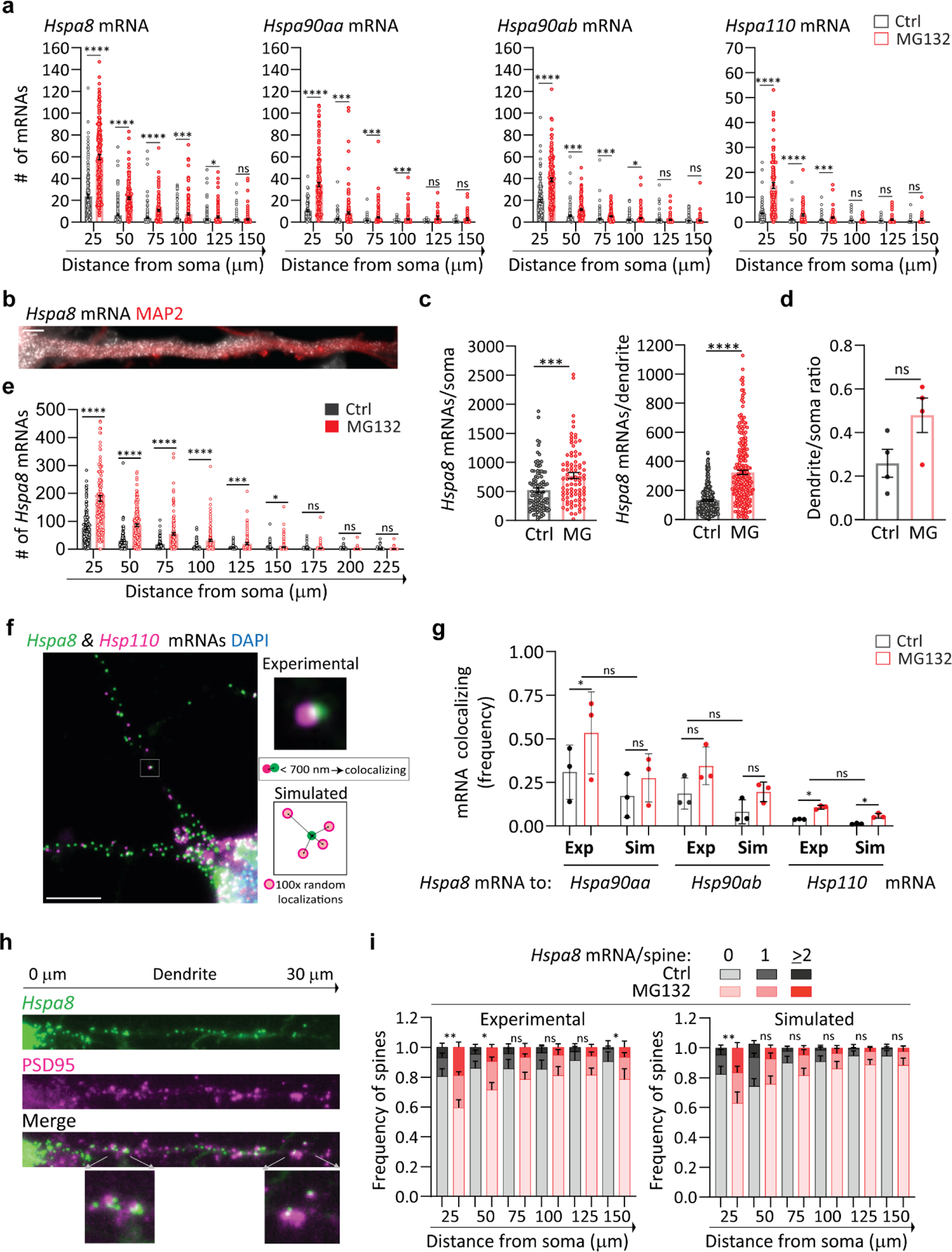
Stress-induced changes in dendritic HSP mRNA localization in primary neurons. **(a)** Quantification of the dendritic mRNAs located in 25-μm bins based on their distance from the soma. Data are the mean ± SEM of three independent experiments (*n* = 88–185 dendrites; dots indicate individual dendrites). **(b)** smFISH detection of *Hspa8* mRNAs in the dendrites of an MG132-stressed primary mouse motor neuron stained with MAP2. Scale bar = 5 μm. **(c)** Quantification of somatic and dendritic *Hspa8* mRNAs in Ctrl and MG132-stressed motor neurons. Data are the mean ± SEM of four independent experiments (*n* = 87–100 neurons and *n* =237-280 dendrites). (**d**) Ratio of *Hspa8* mRNA per area of dendrite or soma (in pixels) in Ctrl and MG132-stressed motor neurons analyzed in C. **(e)** Quantification of *Hspa8* mRNAs per 25-μm bin in experiment **c** (*n =* 154-160 dendrites). **(f)** Two-color smFISH detection of *Hspa8* and *Hsp110* mRNAs in a primary hippocampal neuron stressed with 10 μM MG132 for 7 h. The square shows a magnified view of two colocalization mRNAs and the scheme represents the calculation of colocalization (< 700 nm away) in the experimental data and random simulations. **(g)** Frequency of colocalization between *Hspa8* mRNA and *Hsp90aa*, *Hsp90ab*, or *Hsp110* mRNAs per dendrite in Ctrl and MG132-stressed primary hippocampal neurons. Exp indicates experimental data. Sim indicates simulated data that is the average of 100 random simulations of the positions of each detected *Hspa90aa*, *Hspa90ab*, or *Hsp110* mRNA in a specific dendrite. Data from A. *P* < 0.01; *, *P* < 0.05; ns, not significant (by Wilcoxon *t*-test). **(h)** Detection of *Hspa8* mRNAs in relation to the dendritic spines, identified by anti-PSD95 IF. The distances shown are in relation to the soma. The lower images show magnifications of mRNAs localizing to the dendritic spines in the areas indicated by the arrows. **(i)** Frequency of dendrites with 0, 1, and 2 or more *Hspa8* mRNAs localizing within 600 nm of the center of the PSD95 IF signal in Ctrl and MG132-stressed primary hippocampal neurons from panel a. Dendritic spines were assigned to 25-μm bins based on their distance from the soma. Six experiments were analyzed. Simulated data are the average of 100 random simulations of the positions of each detected *Hspa8* mRNA in the specific dendritic bin. *P* < 0.01; *, *P* < 0.05; ns, not significant (by chi-squared test).

We examined whether mouse primary spinal cord motor neurons, which have more extensive and thicker dendrites than hippocampal neurons, similarly regulate *Hspa8* mRNA. Dissociated cultures of embryonic day 13 mouse spinal cords were matured for at least 3 weeks, when cultured motor neurons express properties of their *in situ* counterparts ^97^. Upon MG132 (10 μM for 7 h) exposure, motor neurons increased *Hspa8* mRNA concentrations in their somas and dendrites, promoting dendritic localization (**Fig. 3b–3d**); however, *Hspa8* mRNA was more concentrated in the proximal dendrites in both stressed and non-stressed neurons (**Fig. 3e**). These results indicate that the regulation of HSP mRNA localization is shared by different neuron types and that their movement towards distal dendrites is constrained or subjected to retrograde transport.

RNAs, proteins, and components of the translation machinery phase separate into neuronal granules of ∼700 nm in diameter with distinctive compositions ^34,37,98^. The concomitant localization and similar dendritic distribution of all HSP mRNAs suggested that they can be assembled and transported in the same neuronal granules. To identify mRNAs traveling with the most abundant dendritic mRNA, *Hspa8*, we performed two-color smFISH in hippocampal neurons to detect *Hsp90aa*, *Hsp90ab*, or *Hsp110* localizing within 700 nm of, and thus coexisting with, each *Hspa8* molecule (**Fig. 3f**). We observed higher frequencies of *Hspa8* coexisting with any of the other HSP mRNAs in MG132-stressed neurons than in non-stressed neurons (**Fig. 3f and 3g**). To differentiate regulated and random mRNA colocalization, we created a new module in ARLIN ^89^ that randomly simulated the positions of *Hsp90aa*, *Hsp90ab*, or *Hsp110* mRNAs over the dendritic shaft. ARLIN averaged the shortest distances between *Hspa8* and the closest HSP mRNA obtained in 100 random simulations (**Fig. 3f and 3g**). Although the coexistence between *Hspa8* with *Hsp90aa*, *Hsp90ab*, or *Hsp110* mRNAs significantly increased upon exposure to 10 μM MG132 for 7 h, the levels were similar between the experimental and simulated data in control and MG132-stressed conditions (**Fig. 3g**). Similar results were obtained when quantifying the coexistence of *Hsp90aa*, *Hsp90ab*, or *Hsp110* mRNAs with Hspa8 mRNA (**Fig. S3a**). Thus we cannot exclude that the coexistence is by chance and we suggest that HSP mRNAs are most probably packaged and transported in individual neuronal granules, as previously reported for other dendritic mRNAs ^99^.

Considering the concentrated localized translation that occurs in the dendritic spines, we hypothesized that they would require relatively more HSPs to prevent aberrant interactions among unfolded proteins. Thus, we used ARLIN to analyze stress-induced changes in the contiguity between HSP mRNAs (detected by smFISH) and the postsynaptic density (detected by PSD95 IF) ^89^ (**Fig. 3h**). Quantifying mRNAs within 600 nm of the center of the PSD95 signal showed that MG132 stress increased the number of spines containing at least one *Hspa8, Hsp90aa*, *Hsp90ab*, or *Hsp110* mRNA (**Fig. 3h and 3i and Fig. S3b**). Because the concentration of HSP mRNAs is higher in the proximal than in the distal dendrites, we used ARLIN to bin the dendritic shafts into 25 μm segments and organized them by distance from the soma to examine changes in the number of HSP mRNAs localizing with the PSD95 signal. The frequency of spines containing HSP mRNAs was higher in the proximal dendrites than in the distal, with up to ∼ 20% and 40% of spines localizing *Hspa8* mRNAs in control and MG132-stressed neurons, respectively (**Fig. 3i**). To determine whether localization near a dendritic spine was regulated or due to increased HSP mRNA density, we used the ARLIN segmentation module to run random simulations of HSP mRNA localizations while maintaining the position of the PSD95 signal. The contingency between all HSP mRNAs and dendritic spines was similar to that in random simulations over the dendritic shafts of control and MG132-stressed neurons, with not significant differences (*P* > 0.05 (chi-squared test)) observed between the experimental and simulated data in each bin meaning that the same outcome could have arisen by chance (**Fig. 3i and Fig. S3b**). Given the density of the cultures, we were unable to quantify mRNAs in the distal dendrites (> 150 μm from the soma) without introducing errors in their assignment to specific dendrites. We conclude that the increased localization of HSP mRNAs in dendrites upon stress increases the number of spines containing them.

### Function and mechanism of HSP mRNA localization in dendrites

We postulated that the localization of HSP mRNAs in dendrites promotes neuronal survival to MG132 stress. Thus, we distinguished live and dead motor neurons in primary mouse spinal cord cultures that were kept under control conditions, subjected to 10 μM MG132 for 7 h, and recovered from stress for 4 h after MG132 washout from the media (**Fig. 4a**). On average, over 80 percent of neuron remained alive with not significant differences among the three groups (**Fig. 4b**). This result indicates that HSP mRNA localization in dendrites is a response mechanism to sublethal doses of prototoxic stress.

**Fig. 4.**
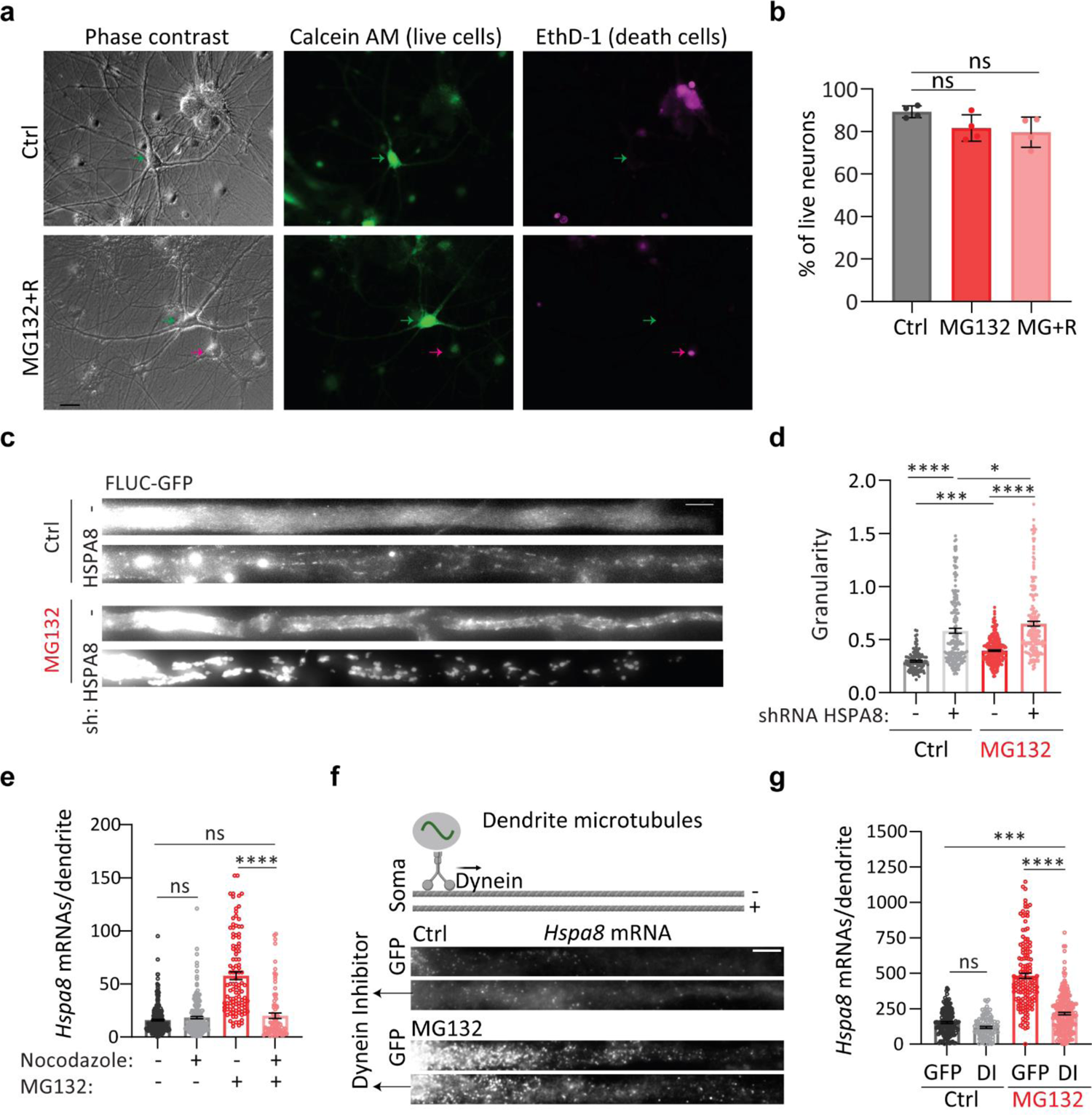
Function and transport of dendritic HSP mRNAs. **(a)** Representative neurons from Ctrl and MG132-stress spinal cord motor neurons double-stained to identify life (Calcein, green) and dead (EthD-1, magenta) neurons, position indicated with arrows. Scale bar = 10 μm. **(b)** Quantification of the life neurons in Ctrl, MG132-stress (10 μM for 7 h), and recovery (10 μM MG132 for 7 h and 4 h after MG132 washout). Three independent experiments (*n* = 8, 99, and 121 neurons). ns; not significant (by Wilcoxon test). **(c)** Representative dendrites from Ctrl and MG132-stressed spinal cord motor neurons expressing the proteostasis reporter plasmid FLUC-GFP with and without HSPA8 knockdown. GFP aggregation is proportional to proteostasis loss. Scale bar = 10 μm. **(d)** Quantification of the GFP signal granularity (the coefficient of variation) in each dendrite in C. Three independent experiments were performed (*n* = 116-334 dendrites from 29-80 neurons). **(e)** Quantification of dendritic *Hspa8* mRNAs in Ctrl neurons and MG132-stressed neurons with and without nocodazole exposure. Data are the median ± SD of three independent experiments. Three independent experiments were performed (*n* = 85–287 dendrites; dots indicate individual dendrite values). **(f)** Schematic of microtubule orientation and dynein transport in dendrites. smFISH detection of *Hspa8* mRNAs in the dendrites of Ctrl and MG132-stressed neurons. Scale bar = 5 μm. **(g)** Quantification of somatic and dendritic *Hspa8* mRNAs in Ctrl and MG132-stressed motor neurons microinjected with a plasmid expressing GFP or a Dynein Inhibitor (DI). Data are the mean ± SEM of three independent experiments (n = 101-2188 dendrites). ****; *P* < 0.0001, ***; *P* < 0.001, **; *P* < 0.01, *; *P* < 0.05, ns; not significant (by 1 way ANOVA).

Considering that *Hspa8* mRNA is the most abundant HSP mRNA in dendrites, we next investigated whether deletion of HSPA8 could weaken dendritic proteostasis as measured using the proteostasis reporter FLUC-GFP ^100^. We first tested the HSPA8 knockdown efficiency of six combinations of two short hairpin (sh)RNAs in Mouse Embryonic Fibroblast (MEFs) and identified a pair that decreased HSPA8 to 7% of its constitutive expression (**Fig. S4a**). We microinjected spinal cord motor neurons with a plasmid encoding the proteostasis reporter FLUC-GFP ^100^. Impaired proteostasis leads to GFP aggregation, which was quantified by the granularity of the GFP signal (coefficient of variation of the GFP signal) ^101^. Four days after microinjections, the granularity of the GFP signal was significantly increased in the dendrites of control neurons depleted of HSPA8 and increased even more upon exposure to MG132 (**Fig. 4c and 4d**). Thus, we conclude that HSPA8 is a major factor in determining neuronal proteostasis in dendrites and its upregulation would be essential to cope with increased protein misfolding upon proteotoxic stress.

We envisioned two non-exclusive mechanisms to promote dendritic *Hspa8* mRNA localization upon stress: active mRNA transport from the soma to the dendrites or enhanced mRNA stability in the dendrites. To distinguish them, we disrupted microtubule polymerization with 0.5 μM of nocodazole to prevent intracellular transport. The significant reduction in the number of dendritic *Hspa8* mRNAs upon combined MG132/nocodazole (10 μM for 7 h/0.5 μM for 11 h) treatment confirmed the importance of active transport from the soma to increase dendritic *Hspa8* mRNA levels upon stress (**Fig. 4e**). As such, longer exposure to MG132, from 1 to 7 h, favored dendritic increases in *Hspa8* mRNA over a somatic increase, and this subcellular distribution was maintained 8 h after MG132 was withdrawn (**Fig. S4b and S4c**). To validate this data and distinguish the roles of transcription and stability, we blocked transcription with actinomycin D (2.5 μg/mL) during MG132-induced proteotoxic stress (10 μM for 7 h) (**Fig. S4d and S4e**). Hippocampal neurons stressed in the presence of actinomycin D lacked smFISH signal for transcription sites, and had fewer HSP mRNAs, with levels similar to those in the control group. The dendritic localization of *Hspa8* mRNA upon stress is significantly lower in neurons exposed to actinomycin D but higher than in the control group (**Fig. S4e**). Thus, mRNA stability plays a minor role in increasing the dendritic concentration of *Hspa8* mRNA upon stress. Overall, these results suggest that stress triggered an initial transport of *Hspa8* mRNAs to dendrites that remained stable during recovery, alluding to RBP-mediated transport.

mRNAs are transported in dendrites bidirectionally by motor proteins: dynein for minus-end and kinesin for plus-end movement along microtubules. To identify the motor protein directing *Hspa8* mRNAs to dendrites, we considered that they harbor microtubules oriented in both plus and minus directions, while axons feature only plus-end-out oriented microtubules (**Fig. 4f**) ^102^. Since dynein is the only motor protein exiting the soma using dendrite-specific (minus-end out) microtubules, we propose it directs HSP mRNAs to dendrites. To test this hypothesis, spinal cord motor neurons were microinjected in the nucleus with a plasmid expressing a dominant negative dynein inhibitor CC1 (DI) fused to GFP (a gift of Dr. Adam Hendricks) ^103^ or GFP alone. CC1 blocks the interaction between Dynein and Dynactin important for dynein-mediated cargo movement in cells. Neurons expressing DI had significantly decreased dendritic *Hspa8* mRNA levels upon MG132 (10 μM for 7 h) (**Fig. 4f and 4g**). However, they still responded to MG132 by increasing the dendritic localization of *Hspa8* mRNA, especially in the most proximal 25 μm of the dendrite (**Fig. S4f**). Thus, targeting of *Hspa8* mRNA to dendrites upon stress depends on microtubule transport partially mediated by dynein.

### HSPs concentrate in the same neuronal compartments as their mRNAs *via* localized translation

Transcript shuttling and local translation is the most efficient way for neurons to target proteins to the dendrites and their spines ^21,22^. We tested whether this regulatory mechanism supports the subcellular distribution of the inducible and constitutive HSP70s (**Fig. 5 and S5**). We quantified the increases in inducible HSP70 (HSPA1A) in the somas and dendrites of spinal cord motor neurons in culture upon MG132 stress (10 μM for 7 h) and after recovery (MG132 washout for 8 h), when the peak of HSPA1A expression was previously reported ^104^. As expected, a significant increase in HSPA1A expression in somas and dendrites was only detected during recovery (**Fig. 5a and 5b**). At this time point, the ratio of dendrites to soma HSPA1A fluorescence signal decreased (**Fig. 5b**), matching the somatic retention of its mRNA (**Fig. 1e and 1h**).

**Fig. 5.**
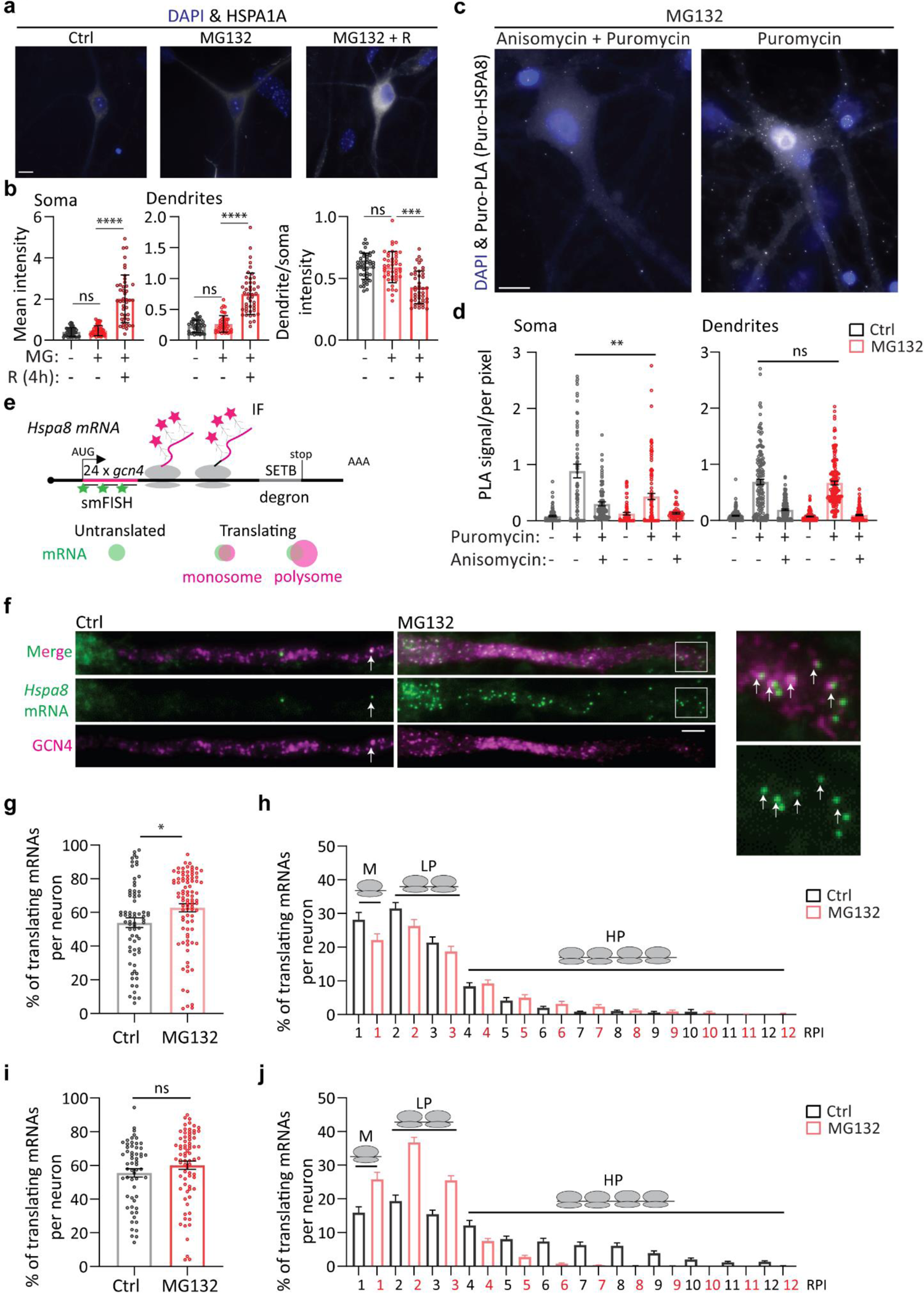
Localized HSP mRNA translation in primary neurons. **(a)** Representative IF detection of HSPA1A in Ctrl, MG132-stressed (10 μM for 7 h), and MG132-recovered (10 μM MG132 for 7 h and washout for 8 h) primary spinal cord motor neurons. **(b)** The plot shows the quantification of HSPA1A in three independent experiments (*n* = 45 total neurons). ****; *P* < 0.0001, ***; *P* < 0.001, ns; not significant (by 1 way ANOVA). **(c)** Representative images of puro-PLA detection of newly synthesized HSPA8 peptides. Scale bar = 10 μm. **(d)** Quantification of the number of puro-PLA signals detected per area of soma and dendrite in three independent experiments (*n* = 51–109 neurons and *n* = 123-163 dendrites; dots indicate individual soma and dendrite values). **; *P* < 0.01, ns; not significant (by 1 way ANOVA). **(e)** Schematic of the *Hspa8* single-molecule translational reporter and the IF-smFISH signals expected for untranslated mRNAs and those being translated by a monosome or polyribosomes. Distention among them was based on the intensity of the IF signal colocalizing with the mRNA, which is proportional to the number of nascent peptides produced from an mRNA. **(f)** Representative IF-smFISH images of dendrites of Ctrl and MG132-stressed primary spinal cord motor neurons expressing the *Hspa8* single-molecule translational reporter. White arrows indicate translating mRNAs. Squares depict the magnified regions. Scale bar = 5 μm. **(g, i)** Quantification of the percentage of *Hspa8* and *Actinb* translated mRNAs per dendrite. Data are the mean ± SD of (g) seven independent experiments (*n* = 70-88 dendrites) and (i) three independent experiments (*n* = 58-74 dendrites). *, *P* < 0.05; ns, not significant (by unpaired *t*-test). **(h, j)** Quantification of the relative nascent peptides intensity (RPI) colocalizing with a translating mRNA from G and I. Media and SEM of translating mRNAs. M = monosomes, LP = light polyribosomes, and HP = heavy polyribosomes.

The *Hspa8* mRNA level increased in both compartments (**Fig. 2c-2f**). Because HSPA8 is a constitutively expressed protein, we used puromycin labeling coupled with Proximity Ligation Assay (puro-PLA) to detect newly synthesized HSPA8 peptides in control and MG132-stressed (10 μM for 7 h) spinal cord motor neurons ^105^. We labeled ongoing peptide synthesis with 5 µM puromycine for 5 min in the presence or absence of the competitor anysomicin (37 µM for 10 min). Specific fluorescent signals for newly synthesized HSPA8 peptides were more abundant in neurons treated with puromycine than in neurons pretreated with anysomicin or DMSO (**Fig. 5c**). Newly synthesized HSPA8 peptides were detected in the soma and dendrites of control and MG132-stressed neurons, suggesting the HSPA8 is locally synthesized in both compartments. Although the number of puro-PLA signals decreased in the soma upon MG132 not significant differences were found in the dendrites (**Fig. 5c and 5d**). These results suggest that localized synthesis determines HSP’s subcellular distribution.

Considering that MG132 decreases global protein synthesis by inducing the integrated stress response ^106^, we investigated whether the efficiency of *Hspa8* mRNA translation changes upon MG132 stress. We generated a single *Hspa8* mRNA translation reporter using the SunTag translation reporter system ^107–110^ (**Fig. 5e**). This plasmid reporter contains *Hspa8*’s 5′ and 3′ untranslated regions (UTRs), coding sequence (CDS), and promoter sequences. We placed 24×GCN4 epitopes at the 5′ end of the CDS to detect nascent peptides as soon as they exit the ribosome tunnel. We added a C-terminal SET binding protein 1 (SETB) degron to decrease the reporter’s half-life and reduce the background.

Translating mRNAs are indicated by colocalization of the smFISH and IF signals detecting the GCN4 nucleotide and amino acid sequences (distance between the centroid of smFISH and IF signal < 300 nm), respectively, while individual signals indicate untranslated mRNAs (green) or fully synthesized proteins that have diffused away from their mRNA (magenta; **Fig. 5e**). We used the IF signal intensity to quantify the relative number of peptides being synthesized per mRNA as a readout of the efficiency of translation of an mRNA and classified them in monosomes (M;1 Relative Peptide Intensity (RPI)), light polyribosomes (LP; 2 and 3 RPI) and heavy polyribosomes (HP; ≥ 4 RPI). The HSPA8 translation reporter resulted in the expression of the expected 125 kDa protein (**Fig. S5a)**. To investigate *Hspa8* translation in neurons, this plasmid was intranuclearly microinjected into mature cultured spinal cord motor neurons. Like endogenous *Hspa8* mRNA, MG132 exposure (10 μM for 7 h) increased the transcription and somatic and dendritic localization of reporter mRNAs (**Fig. 5f and S5b**).

Spinal cord motor neurons were microinjected with the *Hspa8* reporter plasmid, cultures were treated with dimethyl sulfoxide (DMSO; Ctrl) or MG132 (10 μM) 11 h later, and translation was analyzed by smFISH-IF 7 h later (18 h after microinjection). The short window between injection and detection was critical to quantify the efficiency of *Hspa8* mRNA translation in dendrites while avoiding GCN4-HSPA8-SETB accumulation (**Fig. 5f and S5b**). In control spinal cord motor neurons, a few reporter mRNAs localized to the dendrites and were translated by monosomes or polyribosomes throughout the shaft, confirming the reported translation of the endogenous *Hspa8* mRNA by polysome fractionation of neuropil obtained from rat microdissected hippocampi ^111^. Likewise, somatic reporter mRNAs were translated in control and also MG132-stressed neurons. The protein accumulation impaired the accurate quantification of the somatic mRNA translation efficiency, although bright magenta spots depicting polyribosomes colocalized with the mRNA signal in control and MG132-stressed neurons (**Fig. S5b,** magnification in bottom panel). In dendrites, colocalization between the peptide and mRNA signals revealed mRNAs translated by monosomes and polyribosomes at varying distances from the soma (**Fig. 5f**, magnification in bottom right panel). The percentage of mRNAs being translated per dendrite significantly increased upon MG132 exposure (**Fig. 5g**). Similarly, dendritic mRNA translation was more efficient in MG132-stressed neurons, as measured by the shift of mRNAs from the monosome and light polysomes to the heavy polyribosome fraction (**Fig. 5h**). Thus, the constitutive *Hspa8* mRNAs escaped the translational repression associated with MG132 ^112^. These results suggest that *Hspa8* mRNA translation efficiency was increased in response to proteostasis demands.

To determine the specificity in the increased *Hspa8* mRNA translation efficiency, we microinjected spinal cord motor neurons with a plasmid containing the same reporter system, 24xGCN4 epitopes, with the *ActinB* mRNA regulatory elements ^107^. Although the fraction of translating *ActinB* mRNA did not change after stressing the neurons with 10 μM MG132 for 7 h, the mRNAs shifted from the heavy polyribosomes to the light polyribosomes and monosome fractions (**Fig. 5i and 5j**). Thus, while MG132 upregulated *Hspa8* mRNA translation, it suppressed *ActinB* mRNA translation. We also considered that the increased synthesis of HSPA8 could compensate for a high protein turnover and quantified HSPA8 half-life. We co-microinjected a plasmid to express HSPA8 fused to Halo with a GFP-expressing plasmid in primary spinal cord motor neurons. Twenty-four hours after microinjection, neurons were incubated with the photoactive Janilia fluorophore 549 (PA-JF549). GFP-expressing neurons were photoactivated and the decay of the JF549 signal intensity over time was used to calculate the protein half-life to ∼5 days, supporting previous calculations ^106^ (**Fig. S5c and S5d**). These results strongly suggest that combining increased mRNA localization and translation efficiency during stress provides an on-demand source of dendritic HSPA8, that increases in conditions of proteotoxic stress due to the increased load of misfolded proteins.

### The active transport of *Hspa8* mRNAs by FUS ensures dendrite proteostasis

Neurons contain many RBPs relevant to dendritic RNA transport, and several, including TAR DNA binding protein/TARDBP, FUS, and fragile X messenger ribonucleoprotein 1, tightly couple mRNA transport to translation, which is vital for neuronal function ^41–43^. Accordingly, mutations in these RBPs lead to diverse neurodegenerative disorders ^113^. We next identified RBPs that can mediate the dendritic transport of *Hspa8* mRNA by binding to linear or structural zip codes in its 3′ UTR (**Fig. 6a**). We *in vitro* transcribed the *Hspa8* 3′ UTR and LacZ (as a negative control) sequences fused to 2×PP7 stem loops and attached them to amylose magnetic beads using the PP7 capsid protein fused to the maltose binding protein (PCP-MBP) ^86^. Mass spectrometry (MS) identified six RBPs specifically bound to the *Hspa8* 3′ UTR in crude protein extracts obtained from Neuro-2A (N2A) cells kept under control conditions (black *) or exposed to 10 μM MG132 for 7 h (red *) (**Fig. 6a**). Among them, we validated the binding of staufen double-stranded RNA binding protein 2 (STAU2) because of its well-known function in stabilizing and transporting specific mRNAs to dendrites ^31,114,115^ (**Fig. 6b**). We performed two independent pulldowns followed by western blot, and in both cases, the ratio of the large (62 and/or 59 kDa) to small (52 kDa) STAU2 isoforms doubled upon MG132 stress, although its total amount remained constant, which suggest that STAU2 isoforms have differential mRNA affinities upon stress. We also validated the binding of FUS to the *Hspa8* 3′ UTR. The presence of a FUS binding site in the LacZ sequence (ggtgt) probably explained the lack of specificity in the MS profiles (**Fig. 6a and 6b**). FUS was of particular interest because it regulates several steps in mRNA maturation, including transport to dendrites, and *FUS* mutations lead to dendritic retraction in spinal cord motor neurons, leading to ALS and frontotemporal dementia ^27,116–121^.

**Fig. 6.**
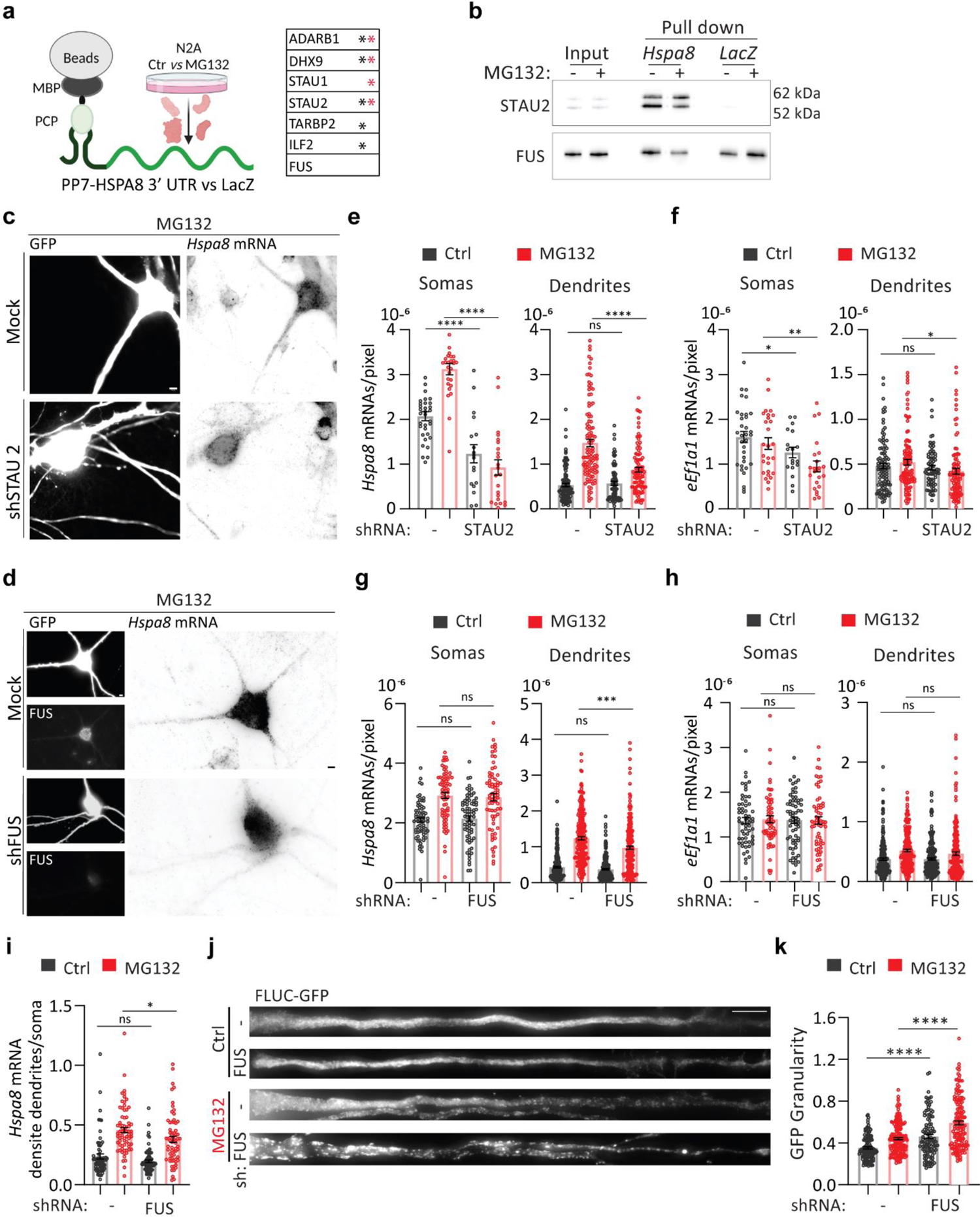
FUS regulates dendritic *Hspa*8 mRNA localization and neuronal proteostasis in mouse motor neurons. **(a)** Schematic of the pulldown strategy used to identify RBPs binding to the *Hspa8* 3′ UTR, and a table of the RBPs identified as specifically bound to the *Hspa8* 3′ UTR in extracts from Ctrl (black *) and MG132-stressed (red *) N2A cells by MS. **(b)** Pulldown experiments to validate the binding of STAU2 and FUS to the *Hspa8* 3′ UTR were analyzed by western blot. I, input; PD, pulldown. **(c, d)** Representative images of primary mouse motor neurons expressing GFP (Ctrl) or GFP and shRNAs against STAU2 (**c**) or FUS (**d**). Three days after microinjection, stress was induced with 10 μM MG132 for 7 h and *Hspa8* mRNA expression was detected by smFISH. Scale bars = 5 μm. **(e– h)** Quantification of the densities of *H*s*pa8* (e**, g**) and *eEf1a1* (**f, h**) mRNAs per pixel of soma or dendrite area in Ctrl and MG132-stressed motor neurons expressing GFP with and without the indicated shRNA expression plasmids. For STAU2, three independent experiments were done ( *n* = 18-33 neurons and *n* = 72-111 dendrites; dots indicate individual soma and dendrite values) and for FUS, five independent experiments were done (*n* = 69-75 neurons and *n* = 224-255 dendrites). ****, *P* < 0.0001; ***, *P* < 0.001; **, *P* < 0.01; *, *P* < 0.05; ns, not significant (by 1 way ANOVA). **(i)** Ratio of the dendrite to soma *Hspa8* mRNA density calculated by averaging the number of mRNAs/pixel of all dendrites of a neuron and dividing it by the number of mRNA/pixel of soma (*n =* 69-75 neurons) from G. *, *P* < 0.05; ns, not significant (by 1 way ANOVA). **(j)** Representative dendrites from Ctrl and MG132-stressed motor neurons expressing the proteostasis reporter plasmid FLUC-GFP with and without FUS knockdown. GFP aggregation is proportional to proteostasis loss. **(k)** Quantification of the GFP signal granularity (the coefficient of variation) in each dendrite in I. Four independent experiments were performed (*n* = 88-138 dendrites). ****, *P* < 0.0001; (by 1 way ANOVA).

To investigate the roles of STAU2 and FUS in the stress-induced dendritic localization of *Hspa8* mRNA, we knocked down their expression in cultured spinal cord motor neurons by co-microinjecting two specific short hairpin (sh)RNAs for each along with a green fluorescent protein (GFP)-expressing plasmid to identify the injected neurons (**Fig. 6c and 6d, and Fig. S6a**). Knocking down all STAU2 isoforms significantly decreased the somatic and dendritic density of *Hspa8* mRNA (quantified as mRNAs per pixel of soma or dendrite area) in control and MG132-stressed neurons (10 μM for 7 h) (**Fig. 6e**); however, MG132 still significantly increased the density of *Hspa8* mRNA in dendrites (*p* < 0.001 by unpaired *t*-test) and preserved the increased in dendritic to somatic *Hspa8* mRNA density in MG132-stressed neurons, demonstrating that STAU2 knockdown did not prevent *Hspa8* mRNA dendritic transport (**Fig. S6b**). To assess specificity, we analyzed the constitutive non-HSP *eEf1a1* mRNA in parallel. STAU2 knockdown had a milder effect on *eEf1a1* mRNA density in the soma and dendrites and did not change the ratio of dendritic to somatic distribution (**Fig. 6f and S6b**). On the contrary, knocking down FUS did not change the somatic concentration of *Hspa8* mRNA but significantly decreased its dendritic density upon MG132 exposure without affecting *eEf1a1* mRNA density in the soma and dendrites (**Fig. 6g and 6h**). As a result, the *Hspa8* mRNA dendrite to soma ratio quantified in single neurons significantly decreased after 7 h exposure to 10 μM MG132 (**Fig. 6i and S6b**). The ratio of dendritic to somatic *eEf1a1* mRNA density did not change upon stress or FUS depletion (**Fig. S6b**). Thus, FUS plays an essential role in targeting *Hspa8* mRNAs to dendrites during stress.

We next evaluated whether FUS knockdown could weaken dendritic proteostasis using the proteostasis reporter FLUC-GFP ^100^. We injected the FLUC-GFP plasmid into spinal cord motor neurons either alone or with the FUS shRNA plasmids. Knocking down FUS increased GFP granularity in dendrites even under control conditions, suggesting that normal FUS levels are essential for neuronal proteostasis under physiological conditions. The loss of dendritic proteostasis upon FUS knockdown was further increased upon MG132 (10 μM for 7 h) treatment, leading to significant increases in GFP granularity and in the sizes of the GFP aggregates (**Fig. 6j and 6k**). These results reveal regulated FUS expression as a determinant of neuronal proteostasis.

### The D290V mutation in HNRNPA2B1 impairs HSPA8 mRNA localization in human-derived motor neurons

Mutations in FUS and other RBPs, including HNRNPA2B1, lead to ALS ^122^. Through a recent collaboration, we found that mouse spinal cord motor neurons microinjected with a plasmid expressing the familial FUS^R521G^ mutation had significantly lower *Hspa8* mRNAs levels in soma and dendrites ^123^. As the *HSPA8* 3′ UTR sequence and length differs between mice and humans, we next investigated whether human motor neurons localize *HSPA8* mRNA to dendrites upon MG132 stress and which RBPs mediate their localization. Interestingly, the human *HSPA8* 3′ UTR sequence contains five putative HNRNPA2B1 binding sites that are not present in mice (**Fig. 7a**). We first validated the specific binding of HNRNPA2B1 to the *in vitro* transcribed human *HSPA8* 3′ UTR sequence fused to 2×PP7 stem loops attached to amylose magnetic beads using PCP-MBP ^86^ (**Fig. 7b**). We performed two independent pulldowns using the crude protein extracts obtained from Ctrl and MG132-stressed HEK293T cells and the LacZ sequence as a negative control. WT HNRNPA2B1 from Ctrl and MG132-stressed cell extracts were specifically bound to the HSPA8 3′ UTR sequence, confirming their physical interaction.

**Fig 7.**
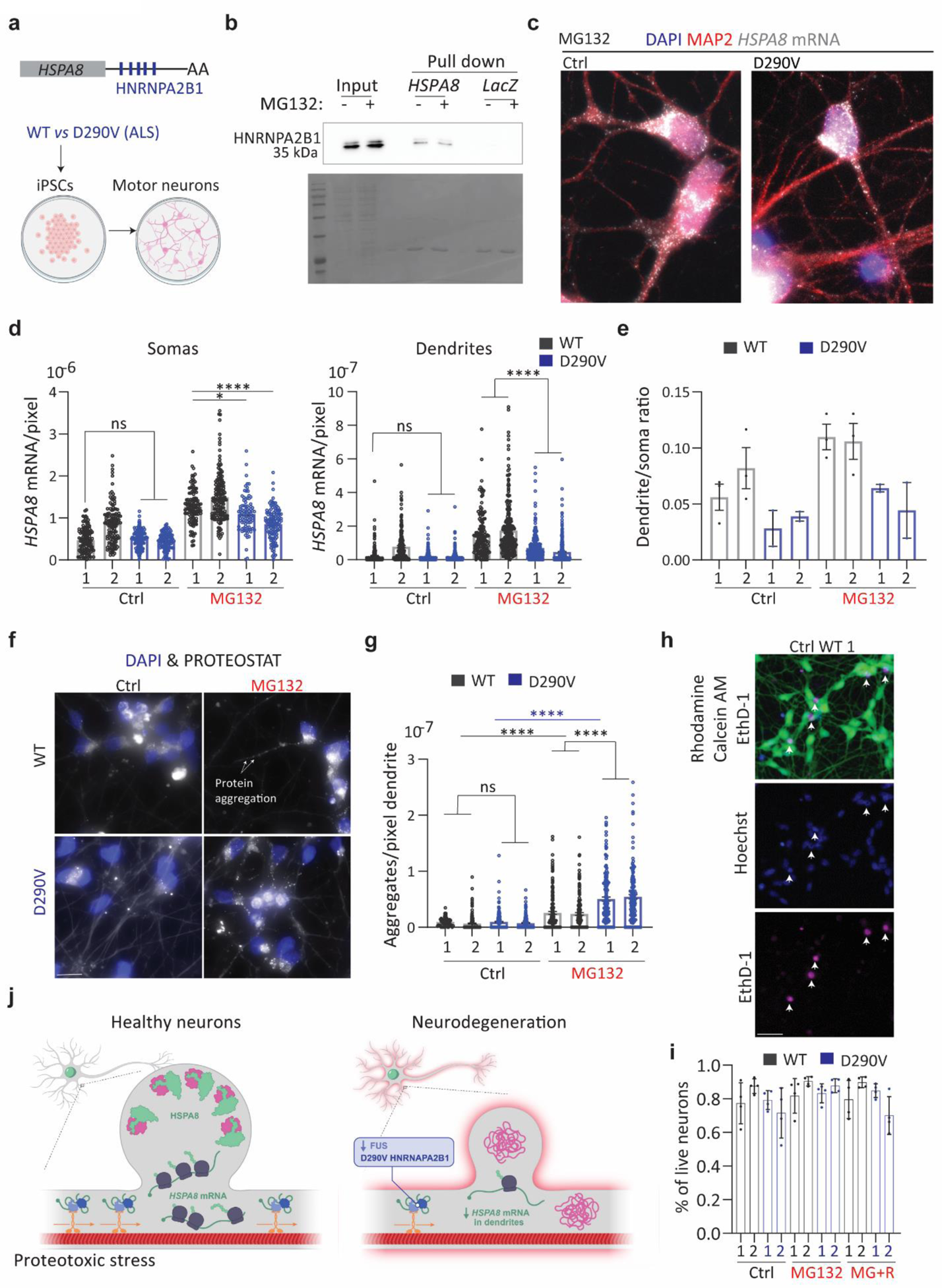
An ALS-associated *HNRAPA2B1* mutation impairs dendritic *HSPA8* mRNA localization in human motor neurons. **(a)** Schematic of human HSPA8 mRNAs and the differentiation of iPSCs from healthy and D190V donors into motor neurons. **(b)** Pulldown experiments to validate the binding of HNRNPA2B1 to the *HSPA8* 3′ UTR were analyzed by western blot. I, input; PD, pulldown; low and high refer to different exposure times of the same blot. **(c)** IF-smFISH to stain dendrites with an anti-MAP2 antibody and detect *HSPA8* mRNAs in MG132-stressed motor neurons differentiated from healthy (WT) donors and patients with ALS carrying the HNRNAPA2B1^D290V^ mutation. Scale bar = 10 μm. **(d)** Quantification of somatic and dendritic *HSPA8* mRNAs in Ctrl and MG132-stressed human-derived motor neurons from the experiments in C. Data are the mean ± SEM of two independent experiments (*n* = 84–155 neurons, n = 150-331 dendrites individual soma and dendrite values indicated by a dot). Motor neurons differentiated from healthy donors WT and patients (P). ***, *P* < 0.001; **, *P* < 0.01; *, *P* < 0.05; ns, not significant (by 1 way ANOVA). **(e)** The ratio of HSPA8 mRNA per pixel of soma or dendrite area in the MG132-stressed motor neurons analyzed in C. **(f)** Representative images of protein aggregates detection in WT and D290V HNRNPA2B derived motor neurons stress with 10 μM MG132 for 7 h. (**g**) Quantification of the aggregates detected by the PROTEOSTAT staining in dendrites in F. Data are the mean ± SEM of three independent experiments (*n* = 42-59 dendrites). ****, *P* < 0.0001; ns, not significant (by 1 way ANOVA). **(h)** Representative neurons from WT MG132-stress iPSCs-derived motor neurons double-stained to identify life (Calcein, green) and dead (EthD-1, magenta) neurons and nuclei (Hoechst, blue). Arrowheads indicate dead neurons in the three channels. Scale bar = 10 μm. **(i)** Quantification of the life neurons in Ctrl, MG132-stress (10 μM for 7h), and recovery (10 μM MG132 for 7h and 4 h after MG132 washout). Three independent experiments (*n* = 83, 99, and 117 neurons). ns; not significant (by Wilcoxon test). **(j)** Summary of conclusions.

In patients with ALS, the D290V mutation in HNRNPA2B1 is rare. However, it promotes the accumulation of the detergent-insoluble HNRNPA2B1 protein in the nucleus and changes the subcellular distribution of mRNAs during stress ^124^. Because of this, human motor neurons differentiated from patient fibroblast-derived induced pluripotent stem cells (iPSCs) do not recover from puromycin stress as well as neurons differentiated from healthy donors ^125,126^. Thus, we compared the ability of healthy (WT) and HNRNPA2B1^D290V^-expressing motor neurons (from two patients each) to localize *HSPA8* mRNAs to the dendrites upon MG132 (10 μM for 7 h) exposure (**Fig. 7c)**.

Importantly, human motor neurons behaved like mouse neurons, increasing the *HSPA8* mRNA level and its dendritic localization upon 7 h exposure to 10 μM MG132 (**Fig. 7c and 7d**). Neurons differentiated from one of the healthy donors (WT2) had significantly higher basal *HSPA8* mRNA levels (by 1-way ANOVA) than the other three lines in somas and dendrites (**Fig. 7d and 7e**). Despite the higher expression, the distribution of *HSPA8* mRNAs in somas and dendrites of Ctrl and MG132-stressed neurons was similar between the two WT-derived motor neurons (**Fig. 7b**). Thus, we compared *HSPA8* mRNA in HNRNPA2B1^D290V^-expressing motor neurons with the healthy WT1. The D290V mutation did not change *HSPA8* mRNA basal expression in soma and dendrites. However, it impaired the somatic accumulation of *HSPA8* mRNAs in HNRNPA2B1^D290V^-derived motor neurons, especially in patient 2. Remarkably, both sets of HNRNPA2B1^D290V^-expressing motor neurons had significantly less dendritic *HSPA8* mRNA than WT motor neurons and this decrease was more significant for patient 2 than patient 1 (**Fig. 7d**). As a result, though both patient cell lines were still responding to MG132 by increasing *HSPA8* mRNA dendritic localization, the distribution ratio of *HSPA8* mRNA in the dendrites relative to the soma was lower in HNRNPA2B1^D290V^ motor neurons than in both WTs upon MG132 stress (**Fig. 7e**). Therefore, HNRNPA2B1’s role in localizing *HSPA8* mRNAs to dendrites is compromised by the D290V mutation.

Considering that *HSPA8* mRNA dendritic localization promotes proteostasis, we next investigated whether HNRNPA2B1^D290V^-derived motor neurons could have a declined proteostasis. We detected the accumulation of protein aggregates with the PROTEOSTAT aggresome kit in Ctrland MG132-stressed iPSCs-derived motor neurons from WT and HNRNPA2B1^D290V^ patients (**Fig. 7f**).

MG132 significantly increased the number of protein aggregates in the dendrites of both WT and D290V neurons; however, the increase was significantly higher with the D290V mutation (**Fig. 7g**). To assess for changes in neuronal survival to stress, we quantified live and dead motor neurons in Ctrl, MG132-stressed (10 μM for 7 h), and stress-recovered (4 h after MG132 washout) cultures (**Fig. 7h**). HNRNPA2B1^D290V^ neurons behaved similarly to WT neurons with an average of 70-95 % of survival under all conditions (**Fig. 7i**). This result confirms that this level of MG132 treatment and the resulting increase in misfolded protein in dendrites was not lethal.

Overall, decreased FUS expression and *HNRNPA2B1* mutation have similar consequences in stressed motor neurons: impaired *Hspa8* mRNA dendritic localization and decreased dendritic proteostasis. Based on our results, we propose a model in which healthy neurons sustain dendritic proteostasis through the regulated localization and translation of HSPs, especially the constitutive chaperone HSPA8, providing an on-demand system to tailor the amounts of HSPs to the load of misfolded proteins. Disruptions in RBPs that impair this localization decrease neuronal proteostatic capacity and prevent synapse formation and transmission, leading to neurodegeneration (**Fig. 7j**).

## Discussion

This study uncovered a novel mechanism to sustain neuronal proteostasis under proteotoxic stress; the partitioning of distinct HSPs in the soma and dendrites through the regulated localization of their encoding mRNAs and subsequent translation. Stress-induced changes in HSP mRNA compartmentalization indicate that distinct proteomes of soma and dendrites are upheld by particular sets of chaperones. It also suggests that after proteotoxic damage the dendritic demands for HSPs exceed the neuron’s capacity to transport individual chaperones from the soma and, instead rely on the competence of an mRNA to produce tens of proteins. The mRNA encoding the constitutive HSP70 HSPA8 is expressed at the highest levels of all HSP mRNAs we detected in dendrites. HSPA8 is central to sustaining proteostasis because of its chaperonin functions in assisting co-translational folding and regulated degradation of proteins by chaperone-mediated autophagy ^127^. HSPA8 also plays moonlighting functions in axonal terminals, where mediates synaptic vesicle fusion and recycling ^64,128^, and in dendrites, where it regulates the shape of dendritic spines ^129^. Besides these critical roles, how HSPA8 localizes to neuronal projections was understudied. Leveraging single mRNA imaging techniques in stressed mouse hippocampal and spinal cord motor neurons in culture has allowed us to discover a specific and regulated *Hspa8* mRNA targeting mechanism to dendrites. Since we did not detect HSP mRNAs in axons, they might localize HSPA8 synthesized in the soma or import from the glia ^130^.

This study identified the post-transcriptional regulation of constitutive *Hspa8* mRNA in neurons, both its boosted dendritic localization and translational efficiency, as a crucial aspect of their stress response. It operates by RBPs recognizing zip codes in the mRNAs, dynein leading them to dendrites where they are transported as individual molecules, and stress-induced translational efficiency. As such, it resembles the induced postsynaptic localization and local translation of the mRNAs encoding an activity-regulated cytoskeleton-associated protein ARC and β-ACTIN that occurs upon synaptic stimulation ^12,25^. We approached uncovering the components directing *Hspa8* mRNAs to dendrites by searching for RBPs that bind to the 3′ UTR sequence of *Hspa8* mRNA since this region is known for its role in mRNA localization. The pool of RBPs identified from Ctrl and MG132-stressed protein extracts was fifty percent identical, suggesting that stress could change the *Hspa8* ribonucleoprotein composition. This might be due to stress-induced changes in the affinity of protein-RNA interactions and/or the stoichiometry of mRNAs and RBPs in the cell, and the unmasking of specific zipcodes. We detected STAU2, well-known for its function in stabilizing and transporting specific mRNAs to dendrites ^31,114,115^. Although STAU2 knockdown in cultured motor neurons reduced the levels of *Hspa8* mRNAs in the soma and dendrites, it failed to prevent the increase in dendritic *Hspa8* mRNAs in response to treatment with MG132. Conversely, FUS was identified as an important player in the subcellular distribution of *Hspa8* mRNA upon stress even though it binds its 3’UTR under Ctrl and MG132-stress conditions. Stress-induced changes in the transcriptome might change the availability of FUS, which can then increase the transport of newly synthesized HSPA8 transcripts to the dendrites. FUS knockdown significantly impaired *Hspa8* mRNA localization in dendrites upon stress but did not completely abrogate it. Thus, FUS might cooperate with additional RBPs binding *Hspa8* 5’ UTR or CDS. It is also possible that the observed effect of FUS to be indirect and operate through the expression regulation of factors involved in the transport of mRNAs.

Besides the fundamental role of FUS in *Hspa8* mRNA transport to dendrites upon stress, we explored this RBP because it is implicated in ALS ^122^, in which dendritic attrition is an early sign of motor neuron damage ^6,131,132^. FUS depletion correlated with a decreased *Hspa8* mRNA dendritic localization and the loss of dendritic proteostasis. Since findings in mouse neurons might not reflect the human situation because of different nucleotide sequences in the *HSPA8* 3′ UTR, we examined iPSC-derived motor neurons. We focused on patient-derived neurons carrying the ALS-linked HNRNPA2B1 mutation D290V because the 3’UTR of the human *HSPA8* mRNA, but not the mouse, has five putative binding sites for HNRNPA2B1. Both lines of HNRNPA2B1^D290V^-derived motor neurons had significantly less dendritic *HSPA8* mRNA than control-derived neurons, indicating that the general principles of RBP regulation of dendritic HSP mRNAs are conserved between mouse and human neurons, with common roles in the neuronal stress response. In addition, the converging actions of different disease forms on *HSPA8* mRNA biogenesis could explain the common loss of proteostasis that characterizes diverse neurodegenerative diseases.

Neurons last the lifespan of an organism, and loss of neuronal proteostasis is a feature of many aging-related neurodegenerative diseases. The localized synthesis of HSPs provides dendrites with the folding resources to aid their protein synthesis and degradation mechanisms in sustaining proteostasis. Perturbations in any of the proteostasis components preclude dendritic homeostasis and lead to dysfunction of neuronal networks and eventually neurodegeneration. As such, boosting HSP transcription has been considered as a therapeutic strategy for neurodegenerative diseases, but has had limited clinical success ^104,133–135^, and has focused particularly on HSPA1A, which is not upregulated in neurons under most conditions ^72,104,133^. Our work stresses the importance of regulating not only the levels of constitutive HSPs, but also the dynamics of their localization to vulnerable regions, such as dendritic spines. These dynamics are crucial to consider when testing potential therapeutics. Sustaining functional synapses is essential for neuronal network functions, and stress on their proteomes would contribute to the impaired connectivity that underlies loss of function early in neurodegenerative disorders, prior to neuronal death ^136–138^.

## Acknowledgments

We would like to thank Dr. Michael Kiebler (Munich Center for Neuroscience) for providing the GFP-STAU2 plasmid and STAU2 antibody, Dr. Adam Hendricks (McGill University) for providing the CC1 expressing plasmid, Dr. Jerry Pelletier (McGill University) for providing the LacZ plasmid, Dr. Lindsay Matthews (McGill University) for MBP-PP7 protein and TEV protease purification, Dr.Tej K. Pandita for the MEFs^HSP70.1-/-^ cell line, Talar Ghadanian and Lokha R. Alagar Boopathy (McGill University) for technical help with the western blots in Fig. S4 and S6, Shruti Iyer and Dr. Adam Hendricks (McGill University) for the HSPA8-Halo and DI plasmids, Dr. Lisa Munter (McGill University) for the hypoxia system, Sethu K. Boopathy Jegathambal and Lokha Ranjani Alagar Boopathy (McGill University) for contributing to the translation program analysis in Fig. 4, and Zoe Wefers and Ryan Huang (McGill University) for help with the quantification in Fig. 2. We thank Dr. Jerry Pelletier for his mentoring and constant support, High-Fidelity Science Communications for manuscript editing, and Margot Riggi for the graphical summary figure.

## Author contributions

Conceptualization: MV, CA, SJT, HD, GWJ. Methodology: CA, JR, PL, SJT, MF, MoV, SX, SM, HD, GWY, and MV. Investigation: CA, PL, SJT, MF, MV, SX, SM, TW, MoV. Visualization and analysis: CA, JR, SJT, MF, MoV, SX, JL, TW. Software development: JR and JL. Funding acquisition and supervision: HD, GWJ, MV. Writing original draft: MV, CA, SX, JR. Revisions: MV, CA, MF, MoV, JR, and JL. Review & editing: MV, HD, PF, GWY, SJT, JR, TW, MoV, MF, and JL.

## Methods

### Neuronal cultures

Mouse primary hippocampal neurons were obtained from postnatal day 0 C57BL/6 and FVB mice and prepared as previously described with small modification ^77,86^. Housing and euthanasia were performed in compliance with the Canadian Council on Animal Care. Isolated hippocampi were kept in Hibernate-A medium and digested with 0.05% trypsin for 15 min at 37 °C. Neurons used in imaging experiments were cultured at low density (50,000 neurons per 35 mm (14 mm glass) dish (MatTek, # P35G-1.5-14-C)) in Neurobasal A Media (Life Technologies, #25300062) supplemented with B-27 (Life Technologies, #A3582801), GlutaMax (Life Technologies, #35050-061), and Primocin (InvivoGen, #ant-pm-1). Neurons used in RNA-seq were cultured at 300,000 neurons per well in 100 μm Transwell membranes (Thermo Fisher Scientific, #35102) in 6-well dishes^80^. Cultures were maintained for at least 2 weeks to ensure neuron maturation and two hundred μl of fresh neuronal media was added every 3 days.

Dissociated spinal cord cultures from E13 CD1 mice were prepared as previously described ^97^. Cells were plated on poly-D-lysine (Sigma, #P6407) - and Matrigel-coated glass coverslips in 6-well dishes. The culture medium was as described ^97^ with the addition of 1% B27 (Gibco Life Technologies, Burlington, ON, Canada, #17504044), 0.032 g/mL sodium bicarbonate, and 0.1 g/mL dextrose.

Cultures were maintained for at least 3 weeks to ensure motor neuron maturation. Human motor neurons were differentiated from iPSCs as previously described ^124,125^. CV-B (wild type) iPSCs were a gift from the Zhang Lab ^139^ and HNRNPA2B1 D290V-1.1 and −1.2 human iPSCs were generated in the Yeo lab ^125^. Human iPSCs were grown on Matrigel-coated 10 cm tissue culture plates.

When cells were 80–90% confluent, they were split into 6-well plates at 1×10^6^ cells/well in 1× N2B27 medium (DMEM/F12+Glutamax, 1:200 N2 supplement, 1:100 B27 supplement, 150 mM ascorbic acid, and 1% Penicillin/Streptomycin) supplemented with 1 μM dorsomorphin (Tocris, #3093), 10 μM SB431542 (Tocris, #1614), 4 μM CHIR99021 (Tocris, #4423) and 10 μM Y-27632 hydrochloride (ROCK inhibitor; Tocris, #1254). The seeding day was counted as day 1. On days 1–5, the cells were refed daily with the same medium as on day 1, but with the ROCK inhibitor reduced to 5 μM. On days 7–17, the cells were refed daily with 1× N2B27 medium supplemented with 1 μM dorsomorphin, 10 μM SB431542, 1.5 μM retinoic acid (Sigma, #R2625), 200 nM Smoothened Agonist, SAG (EMD Millipore, #566660), and 5 μM ROCK inhibitor. On day 18, the cells were either plated on laminin-coated 10 cm plates at 1.2×10^7^ cells per plate for continued differentiation or expanded in motor neuron progenitor MNP medium (1× N2B27 medium supplemented with 3 mM CHIR99021, 2 mM DMH1 (Tocris, #4126), 2 mM SB431542, 0.1 mM retinoic acid, 0.5 mM purmorphamine (Tocris, #4551), and 0.5 mM valproic acid (Tocris, #2815)) on Matrigel-coated plates. To expand motor neuron progenitors, cells were refed every other day with MNP medium. Laminin plates were prepared by serially coating them with 0.001% (0.01 mg/mL) poly-D-lysine (Sigma, #P6407) and poly-L-ornithine (Sigma, #P3655) followed by 20 µg/mL laminin (Life Technologies, #23017015). Cells were refed on day 18 and day 20 with MN medium (1 × N2B27 medium supplemented with 2 ng/mL glial cell-derived neurotrophic factor, 2 ng/mL bone-derived neurotrophic factor, and 2 ng/mL ciliary neurotrophic factor (all from R&D Systems, #212-GD, #248-BD, and #257-CF, respectively) supplemented with 1.5 μM retinoic acid, 200 nM SAG, and either 10 μM ROCK inhibitor on day 18 or 2 μM ROCK inhibitor on day 20. On days 22 and 24, cells were fed with MN medium supplemented with 2 μM DAPT and 2 μM ROCK inhibitor. On day 25, cells were split onto laminin-coated glass coverslips in a 12-well plate at 6.7×10^6^ cells/ well in MN medium supplemented with 10 μM ROCK inhibitor. On day 27, cells were fed with MN medium supplemented with 2 μM ROCK inhibitor. On day 29, cells were stressed with 10 μM MG132 (Sigma, # M7449) for 7 h at 37°C. Cells were then fixed in 4% paraformaldehyde in phosphate-buffered saline and 5 mM MgCl2 (PBSM) for 1 h at room temperature, washed once with 0.1 M glycine in PBSM for 10 min, and stored in PBSM at 4°C for IF staining and mRNA FISH.

### Neuronal manipulation

Neurons were stressed *via* 10 μM MG132 (Sigma, #M7449-1000uL) for the indicated times, hypoxia-reoxygenation (1% O_2_ for 3 h and 4 h recovery at 5% O_2_) using a hypoxia glove box (BioSpherix Xvivo System Model X3), or incubation with oligomers made from 500 nM amyloid-β (1–42) monomers (rPeptide, #1163-1) ^140^. Transcription was inhibited with 2.5 µg/mL Actinomycin D (Sigma, #A9415) for 7 h at 37⁰C. Microtubules were depolymerized with 0.5 µM Nocodazol (Millipore Sigma, # M1404) for 11 h. To quantify the efficiency of *ActinB* mRNA translation, 400 μM Auxin (Sigma, # I3750) was added to the neuronal culture media and the fixation solution. As mature neurons cannot be transfected, plasmids were introduced into primary cultured mouse motor neurons by intranuclear microinjection. The injectate (the plasmid in 50% Tris-ethylenediaminetetraacetic acid (EDTA), pH 7.2) was clarified by centrifugation prior to insertion into 1 mm diameter quick-fill glass capillaries (World Precision Instruments) pulled to fine tips using a Narishige PC-10 puller (Narishige International USA, Inc., NY, USA). Cultures on coverslips were bathed in Eagle’s minimum essential medium without bicarbonate, supplemented with 5 g/L glucose, and adjusted to pH 7.4 in 35 mm culture dishes on the stage of a Zeiss Axiovert 35 microscope (Carl Zeiss Microscopy, LLC, USA) and microinjected using a Transjector 5246 or a FemtoJet Transjector and a Micromanipulator 5171 (all from Eppendorf, Hamburg, Germany). Following microinjection, coverslips were placed in a regular culture medium containing 0.75% Gentamicin (Gibco) and maintained at 37°C in a 5% CO_2_ environment until analysis.

### Plasmid transfection and analysis

Plasmids expressing shRNAs in a lentiviral backbone were obtained through the McGill University library (https://www.sidonghuanglab.com/pooled-screening-libraries/service-request/) (**Reagents)**. They were transfected by calcium phosphate into 293T cells or by Jetprime (VWR, #CA89129-922) MEFs and knockdown efficiency was tested 72 h later by western blotting (**Ragents**).

### IF and smFISH

Detailed protocols for these methods have been previously described for neurons and Mouse Embryonic Fibroblast (MEFs) ^77^. RNA FISH probes were designed using the Stellaris Probe Designer (LGC Biosearch Technologies; masking level: 5, oligo length: 20, minimum spacing: (**Reagents**)).

### Image acquisition and analysis

SmFISH-IF and puro-PLA images were acquired using a Nikon eclipse Ti-2 inverted widefield microscope equipped with a SPECTRA X Light Engine (Lumencor) and an Orca-Fusion Digital CMOS Camera (Hamamatsu) controlled by NIS-Elements Imaging Software. A 60× 1.40 NA oil immersion Plan Apochromat objective lens (Nikon) was used with an xy pixel size of 107.5 nm and a z-step of 200 nm. Chromatic aberrations were measured before imaging using 100 nm TetraSpeck™ Fluorescent Microspheres (Invitrogen, #T14792) and considered in the downstream pipeline.

Single mRNAs, peptides, and postsynaptic densities were identified with the MATLAB version of the FISH-quant (v3) ^87^ and puro-PLA and PROTEOSTAT signals were identified with custom modifications to Big-FISH ^88^. Post-detection analyses of subcellular mRNA distributions in neurons and simulations were performed with the second version of ARLIN ^89^. The code for ARLIN v1.0 and ARLIN v2.0 can be found here: https://github.com/LR-MVU/neuron. See the corresponding documentation for a detailed explanation of ARLIN’s functionalities. Briefly, in ARLIN v2.0, simulations were improved by mimicking the distributions of real mRNAs when selecting simulated mRNAs. To do this, the dendrite was divided into “bins” of 25 μm. The program first counts *x* real mRNAs found in the first bin of the dendrite (*i.e*., 0–25 μm from the soma). Then, the program selects *x* “simulated mRNAs” (*i.e*., randomly selected pixels) from only the first bin of the dendrite. This ensures that the concentration of simulated mRNAs near the soma matches the concentration of real mRNAs but with random distributions within the bin. The program counts the number of real mRNAs found in each bin, then randomly selects that number of pixels within it as “simulated mRNAs”. With this improved simulation, the statistical likelihood of an mRNA localizing to the synapses or to another mRNA can be calculated. statistics are calculated for the localization of mRNAs to synapses or to another mRNA. The simulation is repeated 100 times and the localization statistics are averaged. This provides a more accurate comparison between random and biologically driven colocalization patterns than in the first version of ARLIN.

To quantify translation efficiency, cell and dendrite segmentations were performed manually using the “Define outlines” tool in FISH-quant. The smFISH and peptide spots in the cells were fit to a 3D Gaussian model based on the point spread function and the analysis was run in batch mode. The x, y, and z coordinates of the mRNAs and peptides in cells were exported as tabulated text files (.txt) recording the identity of each cell in each image file analyzed. We designed a Python pipeline to first calculate the median brightness of peptide spots that were not within the translating range (threshold) of an mRNA spot, (200 nm plus the chromatic aberration). This median brightness defined a single peptide spot as these peptides were not concentrated around a translating mRNA. Secondly, the pipeline assigns each peptide to the closest mRNA in the cell. If the distance between an mRNA and its closest peptide exceeded the translating threshold, then we considered it a non-translating mRNA. To remove repeated mRNAs and peptides within the threshold, we selected the brightest (i.e., brightest) peptide signal, and then the closest to a single mRNA. The percentage of translating mRNA was calculated by dividing the number of translating mRNAs by the total number of mRNAs per cell. Additionally, the program would estimate the number of translating peptides in a single spot by comparing the brightness of the spot to the median brightness of a single peptide defined above (https://github.com/LR-MVU/neuron).

We used Fiji (ImageJ 2.14.0, Java 1.9.0_322) to calculate the quantify the fluorescence intensity signal of HSPA1A and granularity of the FLUC-GFP and Proteostat Agreesome signal. Motor neurons were microinjected with 2 ug of the FLUC-GFP-2M plasmid. At 3 or 4 days postinjection, neurons were stressed with 10 µM MG132 for 7 h at 37°C or kept under control conditions. We detected GFP expression by IF and generated a maximum projection for the GFP channel. Then, we outlined a region of interest to define each soma or dendrite in a neuron and measured its area, mean fluorescent signal, and standard deviation (SD). The mean fluorescent signal divided by the area was used to calculate the fluorescent intensity of HSPA1A. The coefficient of signal variation—the SD divided by the mean— was used as a readout for GFP to calculate the granularity of the signal. A similar pipeline was used to identify protein aggregates in human motor neurons and mouse hippocampal neurons. Human motor neurons or mouse hippocampal neurons were stressed with 10 µM MG132 for 7 h at 37°C, MG132 was washed, and the neurons recovered for 4 h at 37⁰C. Neurons were fixed for 15 min on ice with 4% PFA in 1X PBS, 5 mM MgCl_2_, quenched for 10 min on ice with 1 M glycine in 1X PBS, 5 mM MgCl_2_ and stored at 4⁰C in 1X PBS, 5 mM MgCl2. Neurons were permeabilized for 20 min on ice with 0.1% Triton X-100, 1X PBS, 3 mM EDTA pH 8.0. Protein aggregates were detected using a Proteostat Aggresome detection kit. (Enzo Life Sciences, ENZ-51035-0025) following manufacturer instructions.

To localize newly synthesized HSPA8 protein with Puro-PLA, motor neurons were stressed with 10 µM MG132 for 7 h at 37°C. The translation was inhibited for 10 min with 37 µM Anisomycin (Sigma, #A9789) and/or 5 min. with 5 µM Puromycin (Gibco, #A1113803). Neurons were fixed for 15 min on ice with 4% PFA in 1X PBS, 5 mM MgCl2, quenched for 10 min. on ice with 1 M glycine in 1X PBS, 5 mM MgCl2 and stored at 4⁰C in 1X PBS, 5 mM MgCl_2_. Neurons were permeabilized for 10 min on ice with 0.01% Triton X-100, 2 mM VRC, and 1X PBS. PLA was performed using puromycin antibody (Sigma, #MABE343), Hsc70 antibody (Proteintech, #10654-1-AP) Duolink In Situ PLA Probe Anti-Mouse PLUS (Sigma, #DUO92001), Duolink In Situ PLA Probe Anti-Rabbit MINUS (Sigma, #DUO92005), Duolink In Situ Detection Reagent RED (Sigma, #DUO92008) and Duolink In Situ Wash Buffer, Fluorescence (Sigma, #DUO82049) following manufacturer instructions. We quantify mouse and human motor neuron cell viability with the LIVE/DEAD™ Viability/Cytotoxicity Assay Kit (Thermo Fisher Scientific, #L32250). The neuronal media was replaced by a fresh working solution containing 0.25 μM Deep Red Nucleic Acid Stain and 2 μM Calcein Amin in neuronal imaging media (Hibernate A Low Fluorescence, TransetYX, #256) and nucleus of human motor neurons was also stained with Hoechst dye (Thermo Fisher Scientific, #H1399). After 30 min of incubation, human motor neurons were imaged in a Keyence BZ-X with 20x objective NA 0.75 and overlapping of Hoechst and Deep Red Nucleid Acid stain was used to determine the area occupied by death neurons. Mouse cultures on coverslips were transferred to a specialized live imaging chamber (Quick Change Chamber, Warner Instruments, Hamden CT, USA) and placed on the stage of the Zeiss Observer Z1 microscope. Motor neurons were identified by phase contrast, and images were captured using CY2 and CY5 epifluorescence with a 20X objective NA 0.5. Live cells were distinguished by bright green fluorescence labeling, while dead cells, characterized by compromised membranes, exhibited deep red fluorescence with distinct nuclear labeling. After visual examination, mouse spinal cord motor neurons were classified as live cells or dead cells.

To quantify the half-life of HSPA8, we co-injected 2 μg of a GFP expression plasmid and 2 μg of HSPA8-Halo in mouse spinal cord motor neurons cultured in a MatTek dish. Twenty-four hours after injection, neurons were incubated with 5 μM JF-PA549 for 30 min and washed twice for 15 min in neuronal media before adding neuronal imaging media. Once an injected neuron was identified through GFP expression, photostimulation of a region of interest was done for 10 ms with a LUNF 405 nm laser controlled by the Opti-Microscan XY Galvo scanning unit of the Nikon eclipse Ti-2 inverted widefield microscope. A single image was acquired in the 555 nm channel (25 % power, 100 ms exposure) every 30 min for 14 h using the Nikon Perfect Focus system. After subtraction of the neuronal fluorescence background before photostimulation, the fluorescence intensity of neurons was quantified and the decrease in the fluorescence intensity values was used to calculate the half-life of the protein.

### RNA extraction and RNA-seq

After 17 days in culture^141^, primary hippocampal neurons were washed with PBS, and the somas were scraped from the membrane and placed into a tube. Somas were centrifuged for 2 min at 2,000 × *g* and resuspended in 400 µL ice-cold PBS. The somas were divided into two tubes, and 750 µL of Zymo RNA Lysis Buffer (ZymoResearch, #R1013) was added to each. To harvest the neurites, membranes were cut from the Transwell, placed face down in a 6 cm plate containing 750 µL of Zymo RNA Lysis Buffer, and incubated for 15 min on ice while tilting the plate every few minutes. The solution was transferred into an Eppendorf tube. RNA was isolated using a Zyma Quick RNA Miniprep Kit (ZymoResearch, #R1054). RNA library generation and Illumina sequencing was performed by the University of Montreal Genomic Platform. PolyA capture, Nextseq High Output paired-end run (2 × 75 bp, coverage ∼ 50M paired-ends per sample). The raw data have been deposited in the Gene Expression Omnibus under accession number GSE202202.

RNA-seq analysis was performed on usegalaxy.org. Adaptors and reads with a quality below 20 within 4-base sliding windows were removed using Trimmomatic (galaxy version 0.38.0; https://doi.org/10.1093/bioinformatics/btu170). Trimmed single-end reads were aligned to the mouse mm10 genome using STAR (galaxy version 2.7.8a+galaxy0; https://doi.org/10.1093/bioinformatics/bts635) with default parameters, and the number of reads per transcript was determined using featureCounts (galaxy version 2.0.1+galaxy2; https://doi.org/10.1093/bioinformatics/btt656) using default parameters. Differential gene expression was determined using DESeq2 (galaxy version 2.11.40.7+galaxy1; https://doi.org/10.1186/s13059-014-0550-8) using default parameters. Gene ontology analysis to identify biological processes enriched in differentially expressed genes was performed using geneontology.org.

To validate the data by RT-qPCR, 25 ng of RNA isolated from the soma or neurites was reverse transcribed into cDNA using iScript™ Reverse Transcription Supermix (Bio-Rad) following the manufacturer’s instructions. For qPCR, cDNAs were diluted two-fold in water. PCR was performed in 5 μL reactions consisting of 1 μL DNA, 2.5 μL PowerUp SYBR Green Master Mix (Thermo Fisher Scientific), and 0.25 μL of each primer (at 1 μM) on a Viaa 7 Real-Time PCR System (Thermo Fisher Scientific; 45 cycles). Standard curves were generated using a log titration of N2A genomic DNA (50– 0.05 ng) and used to quantify the cDNA. The primers used are listed below.

### Identification of RBPs binding to the mouse and human

*HSPA8 3′ UTR* PP7-HSPA8 and PP7-LacZ RNA were first PCR amplified from N2A genomic DNA or a plasmid (donated by Dr. Jerry Pelletier) using the primers listed below, and then *in vitro* transcribed with a MEGAshortscript™ T7 Transcription Kit (Invitrogen, #AM1354) following the manufacturer’s instructions. After transcription, RNA was treated with 2 units of Turbo DNase and then purified by phenol-chloroform extraction and ethanol precipitation. RNA was resuspended in 10 mM Tris containing 0.2 U/mL RNaseOUT. Small samples were resolved on a 0.5X TBE agarose gel and their A260/A280 ratios were measured using a nanodrop to verify the purity of the RNAs. PP7-HSPA8 and PP7-LacZ RNA were heated for 2 min at 95°C, allowed to cool to room temperature to allow PP7 loops to form, and stored at −80°C.

To prepare crude N2A extracts, ten 10 cm plates of cells were differentiated into the neuronal phenotype for 3 days: one day in DMEM supplemented with 5% fetal bovine serum (FBS) and 20 μM retinoic acid, one day in 2.5% FBS and 20 μM retinoic acid, and one day in 1.25% FBS and 20 μM retinoic acid. To prepare crude 293T extract, ten 10 cm plates were grown in DMEM supplemented with 5% FBS. Half of the N2A and 293T cells were treated with 10 μM MG132 for 7 h at 37°C. Cells were washed once with ice-cold 1× PBS and pelleted by centrifugation. The cell pellets were washed once with 1× PBS and 1 mM phenylmethylsulfonyl fluoride. The supernatant was removed and cells were stored at −80°C. The pellets were thawed on ice, resuspended in three volumes of N2A lysis buffer (50 mM Tris-HCl pH 7.5, 100 mM NaCl, 1 mM MgCl_2_, 0.1 mM CaCl_2_, 1% IGEPAL CA-360, 0.5% deoxycholic acid, 0.1% SDS, 1 mM phenylmethylsulfonyl fluoride, 1 mM dithiothreitol, 1× Complete Protease Inhibitor (Roche), and 100 U/mL RNaseOUT) and incubated on ice for 10 min. Cells were snap-frozen in liquid nitrogen and thawed on ice twice before 10 min centrifugation at max speed. The crude extract (supernatant) was transferred to new tubes and stored at −80°C. Protein concentration was determined by Bradford assay and a small sample of crude extract was run on SDS-PAGE stained with Coomassie Blue to ensure no protein degradation. The same protocol was used for protein extraction in HEK 293T cells.

In 100 μL reaction, 1.5 μM of PP7-HSPA8 3′ UTR or PP7-LacZ RNA were incubated with 2 μM MBP-PP7 in RNA-IP buffer (20 mM Tris pH 7.2, 200 mM NaCl, 1 mM EDTA pH 8.0, 5 mM dithiothreitol, and 0.01% IGEPAL CA-360) for 1 h on ice. Magnetic amylose beads (100 μL) were washed twice with RNA-IP buffer, then rotated with PP7-HSPA8 3′ UTR or PP7-LacZ bound to MBP-PP7 for 1 h at 4°C. The beads were washed twice with RNA-IP buffer and then resuspended in 5 mL RNA-IP buffer supplemented with 0.01 mg/mL tRNA (Sigma, #10109541001) and 5–10 mg N2A crude extract for MS or 2 mg of crude extract for western blots. After rotating the beads and N2A crude extract for 2 h at 4°C, the beads were washed five times with RNA-IP buffer, resuspended in 50 μL RNA-IP buffer and 6 µg of TEV protease, and rotated for 3 h at 4°C. The cleaved PP7 proteins bound to the HSPA8 3′ UTR or LacZ RNA and their interactors was collected and the beads were incubated in fresh RNA-IP buffer containing TEV protease overnight. The elutions were pooled, and the proteins were analyzed by MS as previously described (Proteomics RIMUHC-McGill University)^142^. Proteins with fold change values > 1.5 and P-values < 0.01 compared to in the control sample were considered HSPA8 3′ UTR interactors. Statistics were performed using total spectral count and a T-test analysis. The raw data has been deposited in the ProteomeXchange Project under accession number PXD046036.

## Data availability statement

The raw data for proteomic analysis has been deposited in the ProteomeXchange Project under accession number PXD046036 and for RNA-seq has been deposited in the Gene Expression Omnibus under accession number GSE202202.

## Code availability

Code for imaging analysis is deposited at Github: https://github.com/LR-MVU/neuron.

## SUPPLEMENTARY MATERIALS

**Figure S1.**
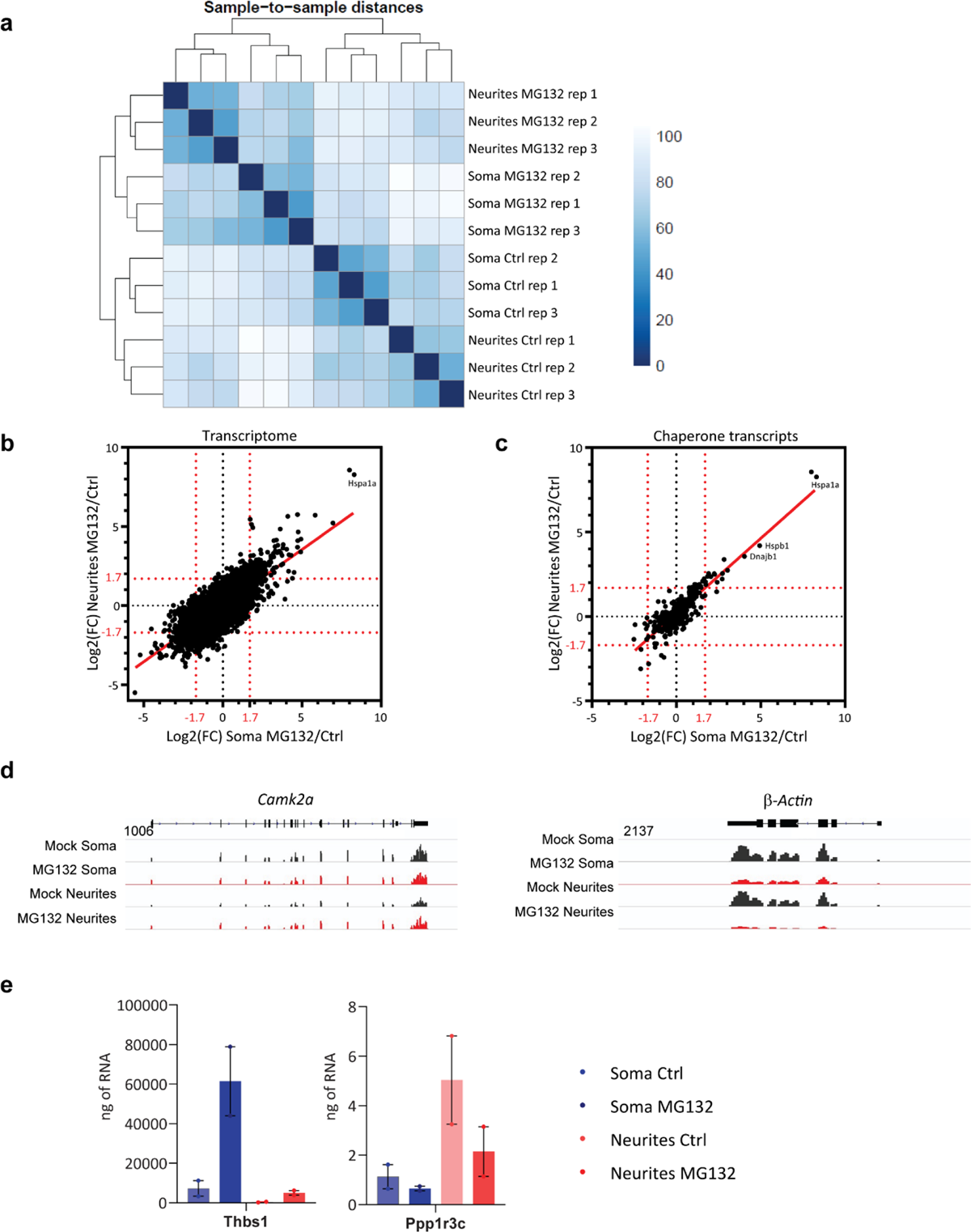
RNA-seq validation. **(a)** Clustering of RNA-seq replicates. **(b, c)** Two-dimensional density plot showing the Log2(FC) of the soma (x axis) and neurites (y axis) in MG132-treated (10 μM for 7 h) versus Ctrl neurons for all genes (B) and molecular chaperone-related genes (C). **(d)** RNA-seq distributions of the *Camk2a* and β-*actin* loci in the soma and neurites of control and MG132-exposed neurons. **(e)** Validation of RNA-seq data by RT-qPCR for the DEGs *Thbs1* (enriched in the soma upon MG132 treatment) and *Ppp1r3c* (enriched in the neurites under control conditions). Data are the mean ± the SEM of two independent experiments.

**Figure S2.**
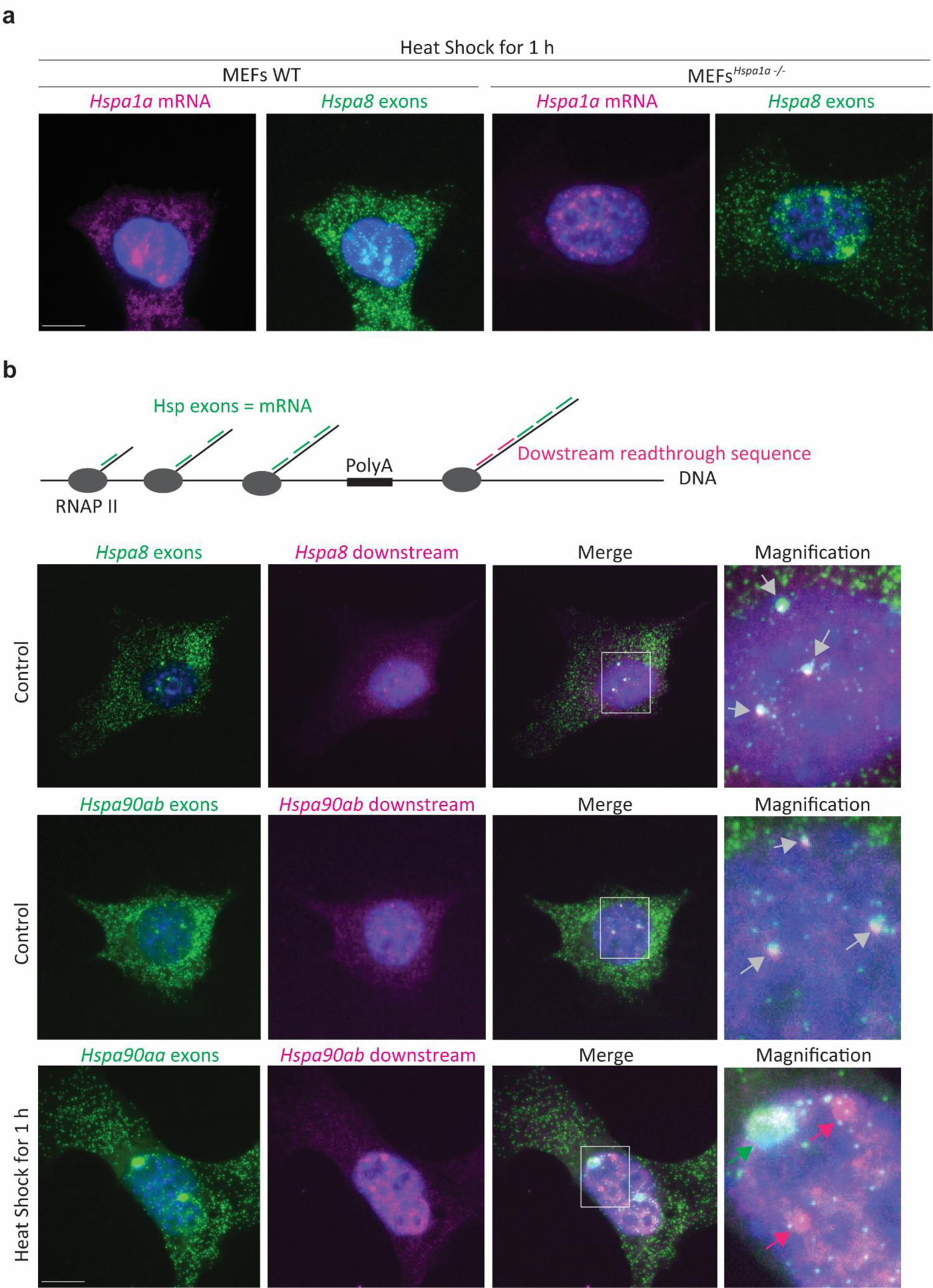
Validation of smFISH probes specificity and transcription site detection. **(a)** SmFISH Detection of *Hspa1a* and *Hspa8* mRNAs in WT MEFs and MEFs^Hspa1a-/-^ subjected to 1 h of heat-shock at 42°C to activate the transcription of Hspa1a. *Hspa1a* mRNA is only detected in the cytoplasm of WT MEFs. **(b)** Upper and middle panels. Identification of *Hspa8* and H*sp90ab* transcription sites in MEFs using two-color smFISH to detect their exon sequences (green) and the genomic region downstream of the PolyA sequence (magenta). White squares indicate magnified images. Gray arrows in magnified regions point to transcription sites where both signals co-localize. Mature mRNAs are only detected with probes for exons. Note that transcription sites are brighter than single mRNAs indicating the presence of more than one transcript per transcription site. Lower panel. Identification of *Hsp90aa* exons (green) and *Hsp90ab* genomic region downstream of the polyA sequence (magenta) in MEFs subjected to 1 h of heat shock to induce *Hspa90aa* transcription. The magnified region shows a lack of co-localization between smFISH probes recognizing different nascent transcripts.

**Figure S3.**
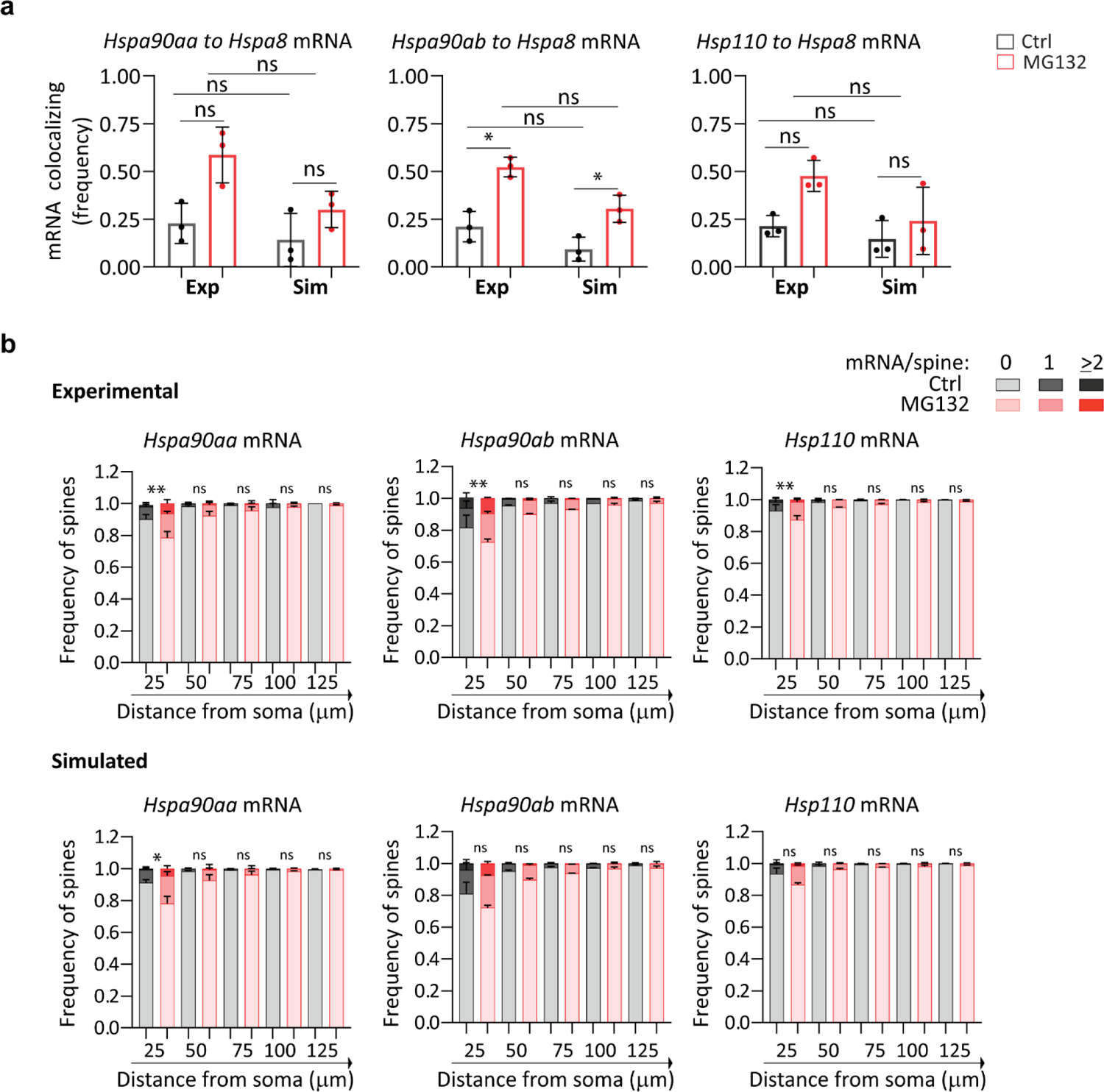
Distribution of HSP mRNAs in the dendritic shaft. **(a)** Frequency of colocalization between *Hsp90aa*, *Hsp90ab*, or *Hsp110* mRNAs and *Hspa8* mRNA per dendrite in Ctrl and MG132-stressed primary hippocampal neurons shown in Fig. 3f and 3g. Exp indicates experimental data. Sim indicates simulated data that is the average of 100 random simulations of the positions of each detected *Hspa8* mRNA in a specific dendrite. Data from Fig. 3h. *P* < 0.01; *, *P* < 0.05; ns, not significant (by Wilcoxon *t*-test). **(b)** Frequency of dendrites with 0, 1, and 2 or more *Hspa90aa*, *Hspa90ab*, or *Hsp110* mRNAs localizing within 600 nm of the center of the PSD95 fluorescent signal in the Ctrl and MG132-stressed primary hippocampal neurons shown in Fig. 3h and 3i. Simulated data are the mean of 100 random simulations of the position of each *Hspa90aa*, *Hspa90ab*, or *Hsp110* mRNAs detected in a specific dendritic bin.

**Figure S4.**
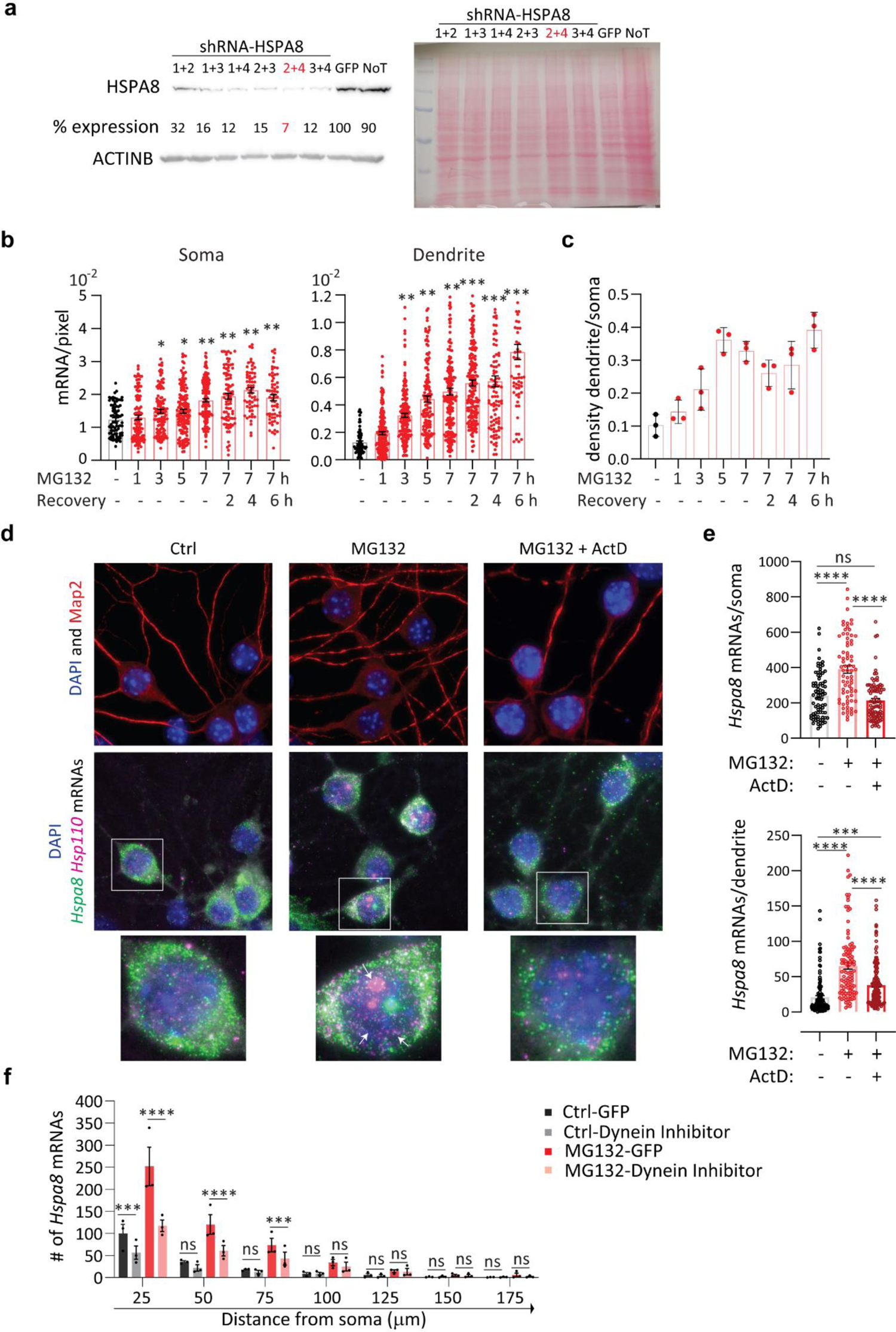
Characterization of the regulated localization of *Hspa8* mRNA in hippocampal neuron dendrites. **(a)** Immunoblot of HSPA8 and β-ACTIN expression in MEFs after 4 days of transfection with the indicated combinations of HSPA8 shRNA plasmids, GFP, and non-transfected (NT). The blot on the right is the ponceau staining of total protein. **(b)** Quantification of the density of *Hspa8* mRNAs per pixel area in the soma or dendrites of Ctrl and MG132-stressed neurons (10 μM) for the indicated times and during recovery from MG132 (Recov) for the indicated times. Data are the mean ± SD of three independent experiments (*n* = 60-155 neurons and *n* = 63-207 dendrites; dots indicate individual soma and dendrite values). **(c)** The ratio of *Hspa8* mRNA per pixel of soma or dendrite area in the Ctrl and MG132-stressed hippocampal neurons analyzed in C. ***, *P* < 0.001; **, *P* < 0.01; *, *P* < 0.05; ns, not significant (by Welch’s *t*-test). **(d)** The upper panels show the detection of the nucleus and dendrites and the lower panel shows two-color smFISH detection of *Hspa8* and *Hsp110* mRNAs of Ctrl, MG132 stressed (10 μM for 7 h), and transcription inhibited (Actinomycin D (ActD) 2.5 µg/mL) MG132 stressed (10 μM for 7 h) primary hippocampal neurons. The square shows a magnified view of a neuron nucleus and arrows point to transcription sites. **(e)** Quantification of somatic and dendritic *Hspa8* mRNAs in Ctrl, MG132-stressed motor neurons transcriptionally active or not. Data are the mean ± SEM of three independent experiments (*n* = 107-134 dendrites; dots indicate individual soma and dendrite values). ****, *P* < 0.0001; ***, *P* < 0.001; ns, not significant (by 1 way ANOVA) **(f)** Quantification of *Hspa8* mRNAs per 25-μm bin in experiment 4F and 4G (*n* = 154-160 dendrites; dots indicate individual dendrite values). ****, *P* < 0.0001; ***, *P* < 0.001, ns, not significant (by *t*-test).

**Figure S5.**
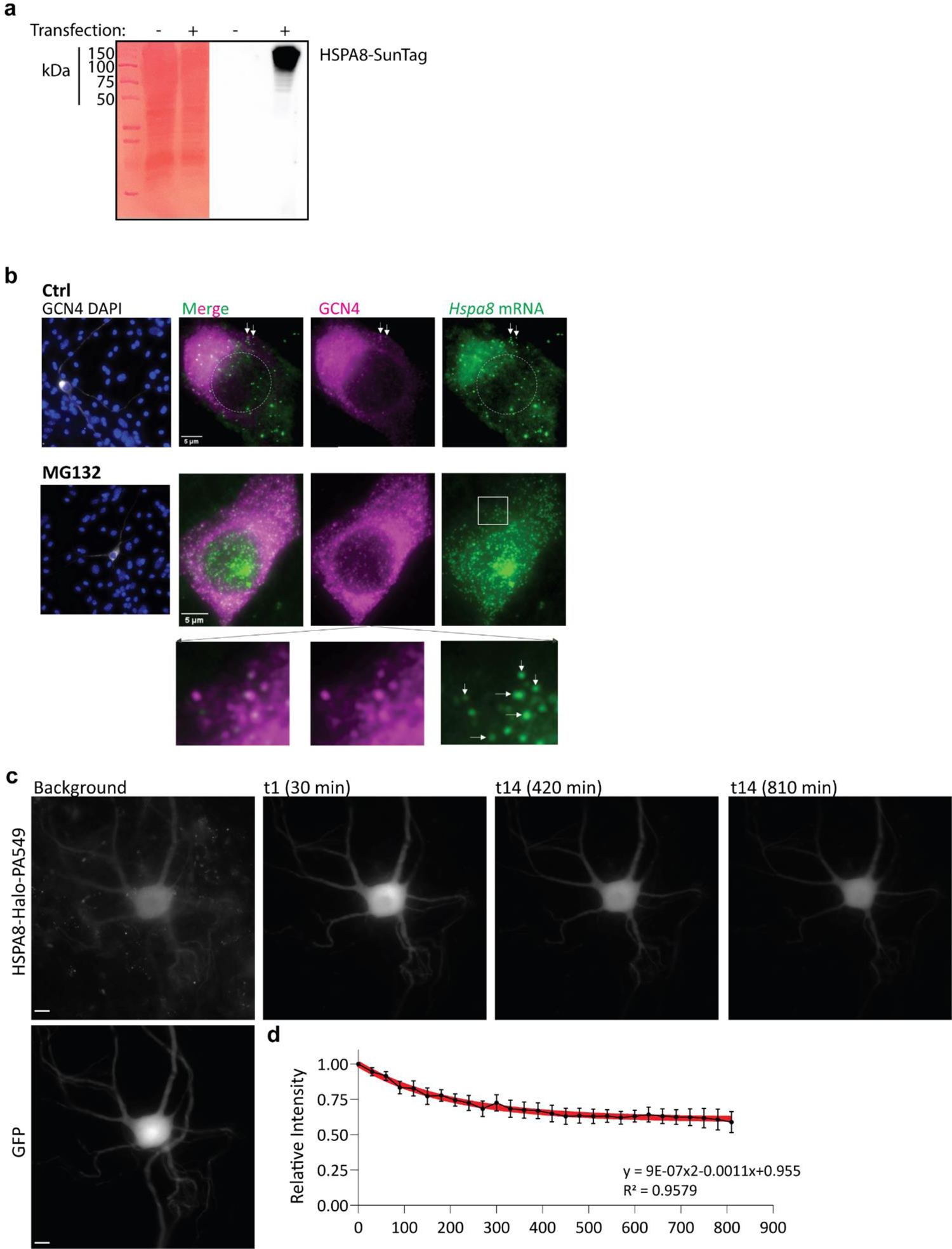
A translation reporter to study localized HSPA8 synthesis in neurons. **(a)** Western blot detection of the HSPA8 single-molecule translational reporter. Left, Ponceau staining of the full protein lysate; right, western blot detecting the reporter’s GCN4 epitopes. **(b)** Representative IF-smFISH images of somas of Ctrl and MG132-stressed primary spinal cord motor neurons expressing the *Hspa8* single-molecule translational reporter. White arrows indicate translating mRNAs. Squares depict the magnified regions. Scale bar = 5 μm **(c)** Representative snapshots of timelapse imaging of primary spinal cord motor neurons co-expressing GFP and HSPA8-Halo. The background is after incubation with PA-JF549 and before photostimulation and t1, t14, and t28 indicate the time imaging acquisition after 10 ms of photostimulation with a 405 nm laser. Scale bar = 10 μm. **(d)** Quantification of PA-JF549 average intensity over time-lapse imaging. The black line and dots are data are the mean ± SEM of 8 spinal cord motor neurons and the red line is the second-order polynomial fit. Equation and R-squared values are displayed.

**Figure S6.**
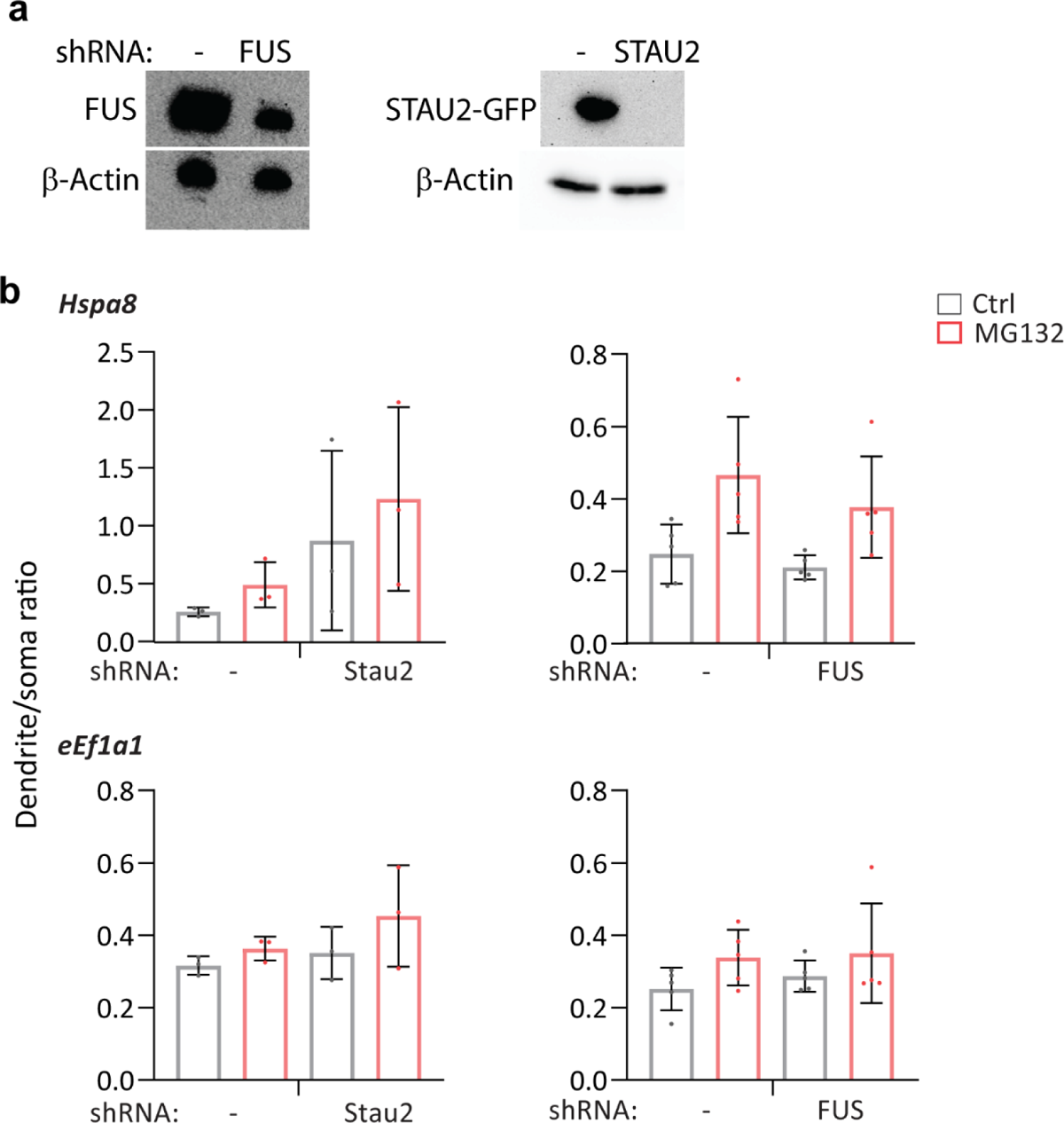
FUS and STAU2 knockdown and HNRNPA2B1 mutation impact the subcellular distribution of *HSPA8* mRNA. **(a)** Western blot detection of FUS and STAU2 in Ctrl and shRNA-transfected 293T cells. β-actin was used as a loading control. **(b)** Ratio of *Hspa8* mRNA per pixel of soma or dendrite area in the Ctrl and MG132-stressed motor neurons analyzed in Fig. 6e–h.

